# AAontology: An ontology of amino acid scales for interpretable machine learning

**DOI:** 10.1101/2023.08.03.551768

**Authors:** Stephan Breimann, Frits Kamp, Harald Steiner, Dmitrij Frishman

## Abstract

Amino acid scales are crucial for protein prediction tasks, many of them being curated in the AAindex database. Despite various clustering attempts to organize them and to better understand their relationships, these approaches lack the fine-grained classification necessary for satisfactory interpretability in many protein prediction problems.

To address this issue, we developed AAontology—a two-level classification for 586 amino acid scales (mainly from AAindex) together with an in-depth analysis of their relations—using bag-of-word-based classification, clustering, and manual refinement over multiple iterations. AAontology organizes physicochemical scales into 8 categories and 67 subcategories, enhancing the interpretability of scale-based machine learning methods in protein bioinformatics. Thereby it enables researchers to gain a deeper biological insight. We anticipate that AAontology will be a building block to link amino acid properties with protein function and dysfunctions as well as aid informed decision-making in mutation analysis or protein drug design.

## 1. Introduction

Amino acids are vital to numerous biological processes, and understanding their physicochemical properties is critical for protein bioinformatics research. The AAindex database [1–4] provides comprehensive quantifications of these properties (*e.g.*, volume, charge, or hydrophobicity) in form of 566 numerical indices (also called scales). Obtained by 149 studies from over six decades of research, these scales constitute valuable features for machine learning models [5–7]. However, the redundancy of AAindex—exemplified by the presence of over 30 scales for hydrophobicity alone— and its sometimes ambiguous annotations impede the development of highly-needed interpretable machine learning models [8,9].

Efforts to cluster the AAindex database have aimed to organize amino acid properties and elucidate their relationships. In 1988, Kenta et al. [1] created the first version of AAindex by collating and hierarchically clustering 222 scales (using agglomerative clustering with single-linkage [10,11]) into four groups: alpha and turn propensity, beta propensity, hydrophobicity, and other properties such as bulkiness. This work was extended by Tomii et al. [2] in 1996, introducing two new groups (amino acid composition and physicochemical properties) for the updated 402 indices, underscoring the importance of studying amino acid scale relationships. An update in 2008 expanded AAindex to 544 scales [4], leading Saha et al. [12] to conduct a consensus fuzzy clustering analysis in 2012 [13], where scales were clustered by various algorithms and then assigned via majority voting, yielding eight clusters. They introduced three new groups: electric properties, residue propensity, and intrinsic properties—while eliminating ‘other properties’. Moreover, they addressed the issue of scales not being clustered into main clusters by assigning them to the ‘intrinsic property’ group. In 2016, Simm et al. [14] clustered 98 hydrophobicity scales, mostly from AAindex, emphasizing their impact on secondary structures. Lastly, in 2018 Forghani and Khani [15] performed a multivariate clustering analysis on the latest version of AAindex (566 scales, 2017), revealing issues with clustering algorithms’ performance and their dependence on settings, particularly for models with pre-defined cluster number such as *k*-means [16]. Using various clustering quality measures, including the silhouette coefficient [17] and the Calinski Harabasz score [18], they determined optimal numbers of clusters: 2, 3, and 9. However, these clusters were not further biologically characterized.

While the existing studies have advanced our understanding of amino acid properties, two limitations hinder their direct use in protein prediction. The first limitation, redundancy, occurs when similar properties are grouped, potentially merging distinct but related properties such as charge and polarity. The second issue, limited interpretability, stems from clustering only based on statistical similarity, which may not reflect the biological meaning or functional relationships. Moreover, current efforts oversimplify the diversity of property scales by confining them to a few clusters without adequately mitigating their complexity. Therefore, a pressing need exists for a more fine-grained and biologically meaningful classification to meet diverse research requirements.

Here, we introduce AAontology, a two-level ontology [19,20] of amino acid scales, a systematic description of scales, and a comprehensive analysis of their relationships. AAontology organizes amino acid property scales into 8 categories and 67 subcategories based on their numerical similarity and physicochemical meaning. It enhances their interpretability, particularly for scale-based machine learning in protein prediction tasks [21–24]. This framework may serve as the foundation for systematically exploring the relationships between physicochemical properties and protein functions (*e.g.*, cellular signaling [25,26], molecular recognition [27], or membrane insertion [28]) and dysfunctions (*e.g.*, oncogenicity [29–31] or aggregation linked to diseases such as Alzheimer’s disease [32–34]). In addition, it may also propel informed decision-making in mutation analysis [35–40] or drug design of proteins such as antibodies [41,42]. Thus, AAontology promises to deepen our understanding of the multifaceted landscape of protein biology.

## 2. Material and methods

### 2.1 Dataset of amino acid scales

We obtained 566 property scales for amino acids from the AAindex database (version 9.2) [43], including their *scale id*, *scale name*, and their one-sentence description, referred to as *scale description*. A further 86 scales related to the solvent accessible surface area (ASA) [44] and hydrophobicity were manually collated from the literature (72 from Lins et al. [44] and 14 from Koehler et al. [45], respectively) because of their general relevance for protein folding [46] and backbone dynamics [47]. After discarding scales due to missing values or complete redundancy, 586 scales remained in our dataset (553 from AAindex, 21 from Lins et al., and 12 from Koehler et al.). For Lins et al. and Koehler at al., we created new *scale ids* by adopting the AAindex naming convention: first authoŕs last name, publication year, and the order of appearance in the publication (LINS030101, …, LINS030121 and KOEH090101, …, KOEH090112). A second publication in the same year is indicated by 02 such as in CHOP780211 (Chou-Fasman, 1978b). Moreover, a *scale name* and a *scale description* were created for each scale using the descriptions provided in the respective publications. Each of the 586 scales was min-max normalized to a [0,1] range (**Supplementary Table 1**).

### 2.2 Representation of amino acid scales

Property scales are represented as arrays **x** containing 20 numerical values, each corresponding to one of the 20 canonical amino acids: 𝐱 = [a_l_, a_2_, … a_20_]. Multiple property scales are represented as an *m* × 20 feature matrix ***X***:

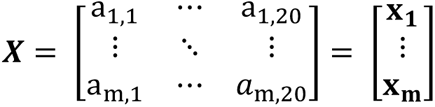

where *m* is the number of property scales.

### 2.3 Representation of scale subcategories by average scales

To compare different sets of scales, we created representative ‘average scales’. Given a predefined set of scales S = {𝐱_𝟏_, 𝐱_𝟐_, … 𝐱_m_}, with *m* denoting the number of scales, an average scale **X̄** was computed as the arithmetic average of scale values for each amino acid across all scales: **X̄** = [ ā_l_, ā_2_, … , ā_20_], where ā_i_ is the arithmetic average value for the *i*-th amino acid over all *m* scales in S.

Such sets of scale can be derived using clustering algorithms. These algorithms generally partition sets of observations (in our case, the 566 property scales) into subsets or clusters by grouping numerically similar observations together. The central point of a cluster (called cluster center or *centroid*) is the arithmetic mean of the scales within that cluster and corresponds to our defined average scale **X̄**. The scale closest to the *centroid* is the *medoid*, which we refer to as ‘medoid scale’ in our study. Both these concepts will be instrumental in subsequent steps.

In this work, we define scale subcategories (indicated in *capital italic*) as sets of ‘similar’ scales. For instance, the *Volume* subcategory includes all volume-related scales, and its average scale is derived from the mean volume scores of the 20 amino acids across all scales. Scale similarity was determined both based on the literature and Pearson correlation analysis, ensuring a minimum correlation of 0.3 within each subcategory. For example, although hydrophobicity and hydrophilicity scales both reflect polarity concepts, they were kept separate to maintain numerical consistency. This approach lessens the cancellation of effects due to anti-correlated scales, allowing the average scales to provide a more robust consensus on certain properties drawn from various studies.

### 2.4 Scale classification and naming by automatic assignment and manual refinement

Leveraging our AAclust framework [48] as well as heuristic and knowledge-based criteria, we classified the 586 amino acid scales obtained from 151 studies curated in AAindex and two further studies (Lins et al. [44] and Koehler et al. [45]). We defined the following 8 categories (color-coded) based on previous results from the literature [1,2,12,14,15]: ‘ASA/Volume’ (blue), ‘Composition’ (orange), ‘Conformation’ (green), ‘Energy’ (red), ‘Others’ (gray), ‘Polarity’ (yellow), ‘Shape’ (light blue), and ‘Structure-Activity’ (brown). To increase the applicability and interpretability of our classification, we further aimed to subdivide each category into subcategories, each with an average size of 5 to 10 scales, ideally yielding a manageable number of between 50 and 100 subcategories in total.

To this end, we performed the following three steps (**Fig. 1**): scale assignment to categories using bag-of-words (a common technique in text classification [49,50]), automatic assignment of scales to subcategories through AAclust, and manual refinement of the scale assignment to subcategories and categories. To improve the clustering quality, we performed these steps multiple times with different clustering algorithms and AAclust settings, similar to an ensemble clustering approach [51]. The process is shown in **Fig. 2**, with comprehensive details provided in **Supplementary Table 2**.

**Fig. 1.**
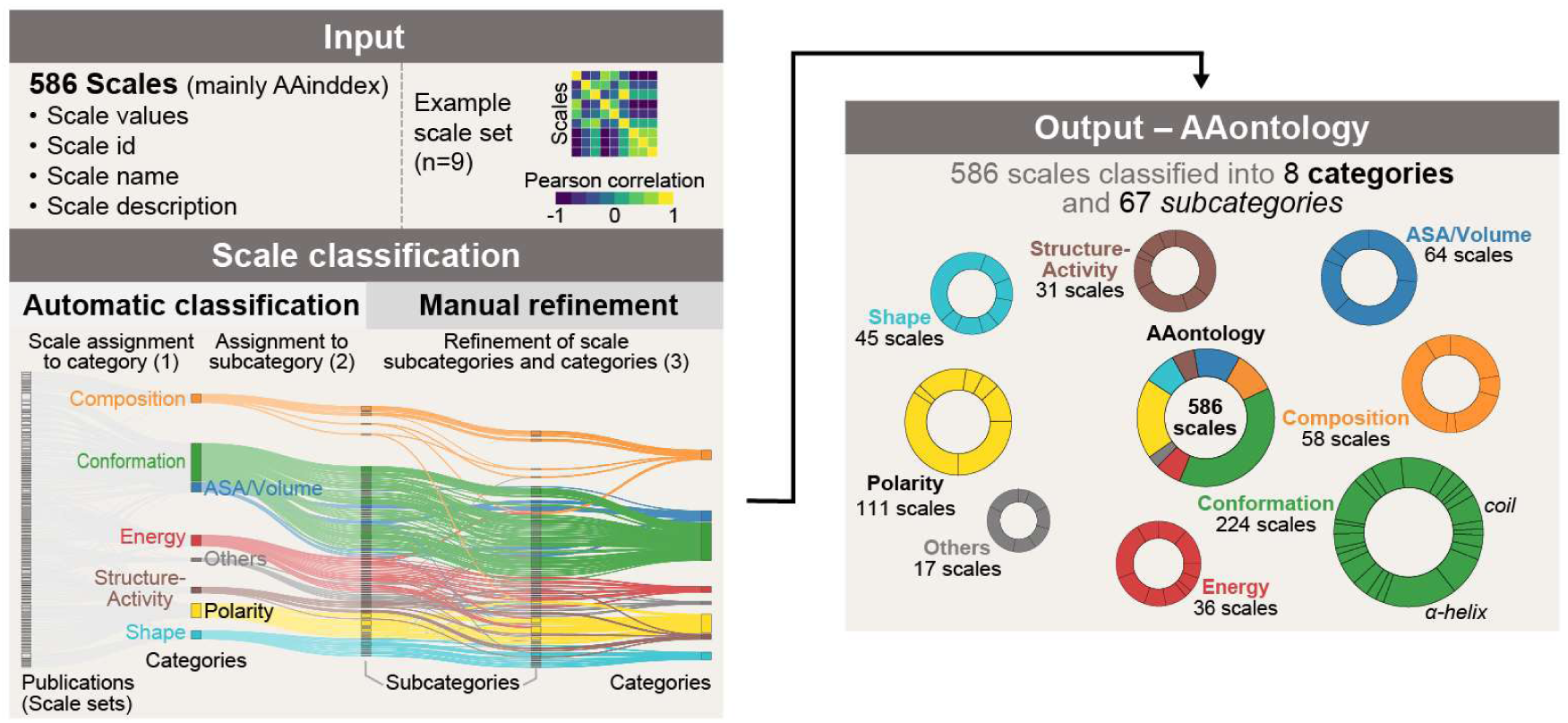
Workflow for creation of AAontology. 586 property scales are automatically classified and manually refined based on their amino acid values, *scale name*, and *scale description*, yielding a two-level classification of 8 categories (color-coded) and 67 subcategories, called AAontology.

**Fig. 2.**
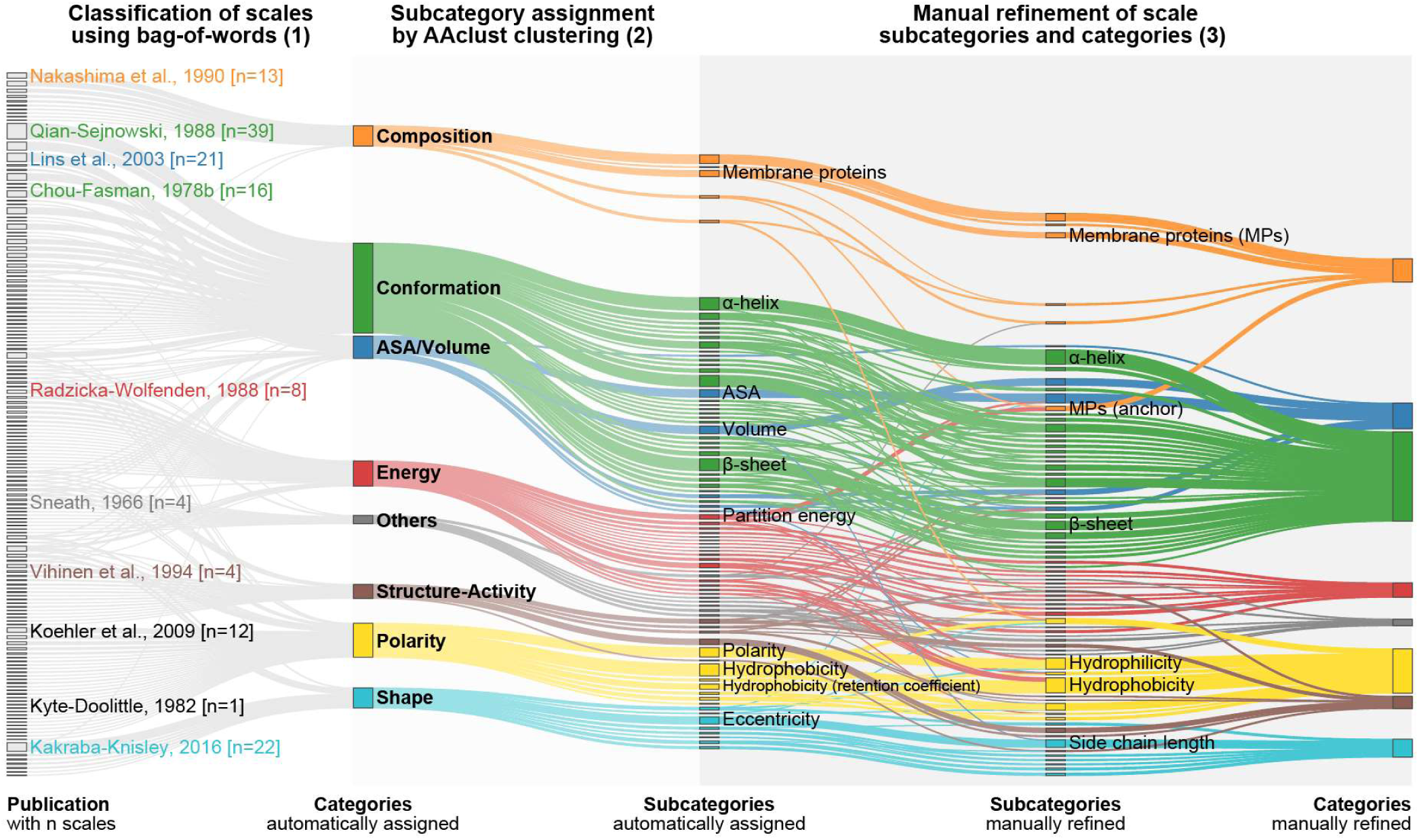
Creation of AAontology. Sankey diagram showing the process of scale classification. 586 amino acid scales from 151 publications were first automatically assigned to 8 different categories using a bag-of-word approach based on their scale description. Key references with the number of containing scales are given. Next, an automatic subcategory assignment was performed using clustering with AAclust [48]. Finally, subcategory and category assignment were manually refined including the renaming of the automatically created subcategories. This procedure was repeated over multiple iterations to refine subcategory names and scale assignment.

#### Classification of scales using bag-of-words

In the field of natural language processing, bags-of-words are sets of terms associated with the same class and they are employed as a simple tool for text classification [49,50]. We defined a bag-of-words for each of the 8 categories: ASA/Volume ({‘accessible surface‘, ‘volume’, …}), Composition ({‘amino acid distribution’, ‘membrane-propensity’, …}), Conformation ({‘β-sheet’, ‘helix’, …}), Energy ({‘charge’, ‘entropy’, …}), Others ({‘mutability’, ‘principal component’, …}), Polarity ({‘hydrophobicity’, ‘polarity’, …}), Shape ({‘graph’, ‘steric parameter’, …}), and Structure-Activity ({‘flexibility’, ‘side chain interaction’, …}). The complete bag-of-words for each category are given in **Table 1**. For each scale and category, the occurrence of these words in the *scale name* and the *scale description* was counted, referred to as word count, and the scale was assigned to the category for which it had the highest word count.

**Table 1.** Scale categories with their bag-of-words and the number of respective subcategories before and after manual refinement (indicated in bold).

| Category | Bag-of-words | # subcategories |
| --- | --- | --- |
| ASA/Volume | ‘accessibility’, ‘accessible surface’, ‘buriability’, ‘buried’, ‘bulkiness’, ‘exposed residue’, ‘size’, ‘van der waals’<sup>#1</sup>, ‘volume’, ‘weight’ | 5<br>**5** |
| Composition | ‘amino acid distribution’, ‘composition’, ‘frequency of occurrence’, ‘membrane preference’, ‘membrane proteins’, ‘mesophilic proteins’, ‘proteins of mesophiles’, ‘mt proteins’, ‘multi spanning proteins’, | 4 (+ 1 unclassified)<br>**5 (+ 1 unclassified)** |
|  | ‘nuclear proteins’, ‘proteins of thermophiles’, ‘proteins of mesophiles’, ‘sequence frequency’, ‘single spanning proteins’, ‘thermophilic proteins’, ‘transmembrane regions’ |  |
| **Conformation** | ‘alpha-helix’, ‘aperiodic indices’, ‘average relative fractional occurrence’, ‘bend’, ‘beta-sheet’, ‘beta-strand’, ‘beta-structure’, ‘coil’, ‘conformational state’, ‘extended’, ‘helical’, ‘helix’, ‘linker’, ‘loop’, ‘normalized frequency’, ‘pi-helices’, ‘pleated-sheet’, ‘relative preference value’, ‘turn’, ‘chain reversal’ | 27 (+1 unclassified)<br>**24 (+1 unclassified)** |
| **Energy** | ‘charge’, ‘electrical effect’, ‘electron-ion interaction potential’, ‘energy transfer’, ‘entropy’, ‘free energies’, ‘free energy’, ‘gibbs energy’, ‘heat capacity’, ‘hydrogen bond donors’, ‘isoelectric point’<sup>#2</sup>, ‘non-bonded energy’, ‘nonbonding orbitals’, ‘partition coefficient’, ‘partition energies’, ‘partition energy’, ‘polarizability’, ‘transfer energy’ | 15 (+1 unclassified)<br>**9 (+1 unclassified)** |
| **Polarity** | ‘amphiphilicity’, ‘cornette et al’*, ‘hydration’, ‘hydrophobic moment’, ‘hydrophobic parameter’, ‘hydrophobicity’, ‘hydrophilicity’, ‘hydropathy’, ‘pK’, ‘polar requirement’, ‘polarity’, ‘refractivity’, ‘retention coefficient’, ‘surrounding residues’ | 5 (+1 unclassified)<br>**6 (+1 unclassified)** |
| **Shape** | eccentricity’, ‘eigenvalue’, ‘graph’, ‘kakraba-knisley’*, ‘longest chain’, ‘prabhakaran-ponnuswamy’*, ‘reduced distance’, ‘side chain’, ‘steric parameter’, ‘value of theta’ | 6 (+1 unclassified)<br>**6 (+1 unclassified)** |
| **Structure-Activity** | ‘average interactions’, ‘b values’, ‘chemical shift’, ‘contact number’, ‘equilibrium constant’, ‘flexibility’, ‘interactivity’, ‘side chain interaction’, ‘signal sequence’, ‘site occupied by water’, ‘spin-spin coupling’, ‘stability’, ‘zimm-bragg parameter’ | 4 (+1 unclassified)<br>**6 (+1 unclassified)** |
| **Others** | ‘bitterness’, ‘kerr-constant’, ‘melting point’<sup>#3</sup>, ‘mutability’, ‘optical rotation’, ‘principal component’, ‘principal property’, ‘rf rank’ | 7 (+1 unclassified)<br>**6 (+1 unclassified)** |

<sup>#1</sup> ‘van der waals’ refers not to van der Waals forces but rather to van der Waals volume or radius, properties related to ‘ASA/Volume’.
<sup>#2</sup> ‘isoelectric point’ is assigned to ‘Energy’ since it is related to the net charge of residues.
<sup>#3</sup> ‘melting point’ is attributed to ‘Others’ due to ambiguous assignments with other categories—it could be associated, for example, with ‘Energy’ (*i.e.*, as a measure of heat capacity) or to ‘Structure-Activity’ (*i.e.*, as a measure of stability).
\* Names of authors contributing various scales to a specific category.

525 out of 586 scales were classified by assigning a scale to a scale category when at least one word within the categorýs bag-of-words occurred in the *scale name* or the *scale description*. If scales were assigned to multiple categories, the category with fewer scales was preferred to achieve a more balanced partition of scales. The remaining 61 scales were classified manually using additional information from the literature.

#### Subcategory assignment by AAclust clustering

Leveraging our AAclust framework [48], we subdivided the categories hierarchically into subcategories, each named according to term counts from *scale names*, such as the *Entropy* subordinated to the ‘Energy’ category.

In the first step, scales within each category were clustered . We used AAclust in conjunction with *k*-means [16] and agglomerative clustering [10] (with average, complete, ward, single linkage methods [11]). These clustering models demonstrated excellent performance in our previous AAclust study, and were used as implemented in the scikit-learn library [52]. For AAclust, we set a minimum within-cluster Pearson correlation of 0.3, ensuring a positive correlation among all scales within each subcategory. Depending on the AAclust merging parameters, the number of clusters varied between 50 and 270 clusters. Cluster models were subsequently evaluated based on the silhouette coefficient [53], Calinski Harabasz score [17], and Bayesian information criterion (BIC) [53], as outlined in AAclust. To obtain a manageable number of 50–100 subcategories, we considered for further steps only the top-performing models that yielded a maximum of 100 clusters over all categories. This was achieved by employing cluster merging in the AAclust approach, a process that reduces cluster numbers by reassigning scales from smaller to larger clusters.

In the second step, we assigned names to each cluster by analyzing term counts in *scale names*, thereby naming subcategories. Term lists from each *scale name* were generated, considering both the *scale name* and whole-word substrings (without parentheses). For example, for the *scale names* ‘α-helix’ and ‘α-helix (N-terminal)’, the term lists would be [‘α-helix’] and [‘α-helix (N-terminal)’, ‘α-helix’, ‘N-terminal’], respectively. Term lists were then combined, and term counts within each subcategory were calculated. Additionally, we used AAclust to identify the medoid scale, having the highest correlation to the cluster center. The most frequently occurring term became the subcategory name, such as *Hydrophobicity* occurring in almost all hydrophobicity scales. Names were assigned in descending order of cluster size, and if two terms occurred equally frequently the medoid scalés name or the shorter was preferred. Clusters with only one scale or a duplicate name were labeled ‘unclassified’ and merged into a single subcategory within their category, denoted as ‘unclassified Category name’. We excluded five charge-related scales from the clustering process due to their binary nature and later assigned them to distinct subcategories.

This procedure was repeated multiple times with varying models and settings to improve naming consistency and clustering robustness [51]. This iterative process also allowed for the refinement of *scale names* based on subcategory classification, *scale description*, and literature review. For instance, scales such as the ‘TOFT index’ or ‘PRIFT index’ led to the subcategory of *Amphiphilicity (α-helix)*.

Overall, we obtained 7 ‘unclassified’ and 73 meaningful subcategories (**Supplementary Table 2**). The subcategory names largely overlapped with the bag-of-words used for category classification, offering a convenient link between the scales and their respective subcategories.

#### Manual refinement of scale subcategories and categories

We reduced the 73 meaningful subcategories onto 67 subcategories by manual refinement using heuristic and knowledge-based criteria. In this step, for each scale one out of the following four different actions were performed:

- **No changing**: The scale classification is kept, as automatically derived using AAclust.
- **Changing category**: Change of the AAclust classification for the scale category and subcategory.
- **Changing subcategory**: Change of the AAclust classification for the scale subcategory.
- **Renaming subcategory**: Change of the semi-automatically derived subcategory name.

We manually renamed subcategories of 43 scales for clarity and brevity, applying heuristics including shortening (*e.g.*, ‘Principal Component 1’ to *PC1*), summarizing (*e.g.*, ‘Charge donor’ included into *Charge*), deleting unclear names (*e.g.*, ‘Kerr-constant’), and aligning with naming conventions. Notably, we adhered to a general convention for the N- or C-terminal preference in conformational subcategories, as seen in the renaming of ‘β-turn (3rd residue)’ to *β-turn (C-term)*.

Scales were allotted to ‘unclassified’ subcategories when they had a Pearson correlation lower than 0.3 with any scale within their original subcategory and failed to attain a minimum within-cluster correlation of 0.3 with any other subcategory. Further, a scale was deemed ‘unclassified’ when a conclusive literature-based assignment failed. Consequently, 42 of the total 586 scales could not be classified.

Scales were reassigned to different subcategories and categories to enhance the overall Pearson correlation within each subcategory and align better with existing literature. In some cases, scales were allocated to larger subcategories, resulting in a slightly lower minimum within-cluster Pearson correlation, but maintaining a high degree of specificity and consistency in smaller subcategories. For example, despite a higher minimum correlation with the *PC5* subcategory, the ’Relative mutability’ scale was placed into the broader *Mutability* subcategory.

Overall, we manually refined 70% of the automatically assigned scales using these heuristic and knowledge-based criteria, classifying 544 out of the 586 scales into 8 categories and 67 distinct subcategories. Scales within the same subcategory often displayed a high positive correlation (>0.9) or at least a minimum within Pearson correlation of 0.3 (**Fig. 3**).

**Fig. 3.**
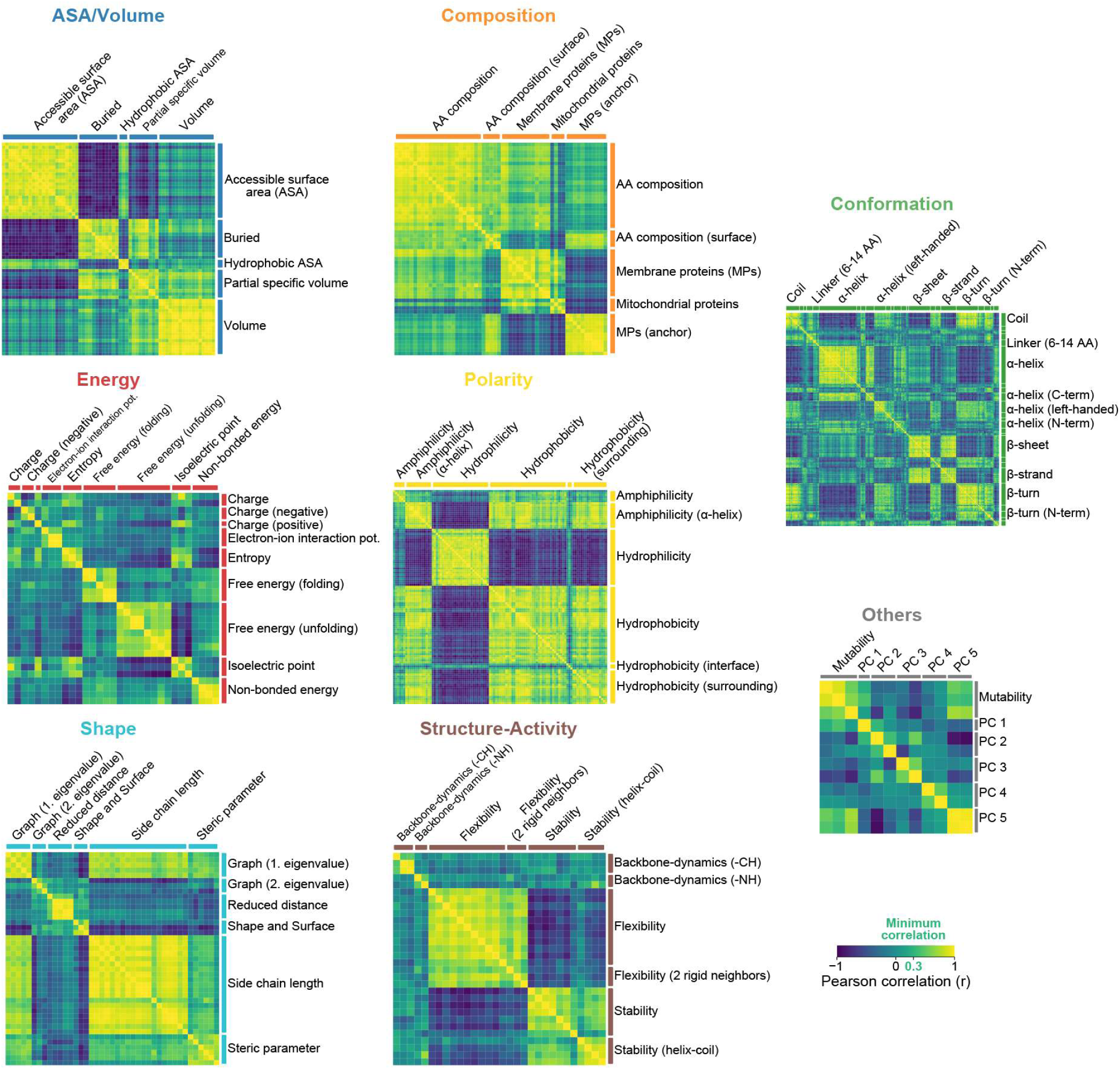
Correlations between scales in each category. Heatmaps for the eight scale categories (color-coded) comprising distinct subcategories with highly positive correlated scales. Each subcategory fulfills a minimum within Pearson correlation of 0.3, which was used as minimum quality criteria for the semi-automatic scale classification.

## 3. Results

The two-level AAontology classification of 586 amino acids scales (from AAindex and two further studies) into 8 categories and 67 subcategories is depicted in **Fig. 4**. This taxonomic hierarchy [19,20] provides meaningful relationships at the level of individual scales and scale subcategories.

**Fig. 4.**
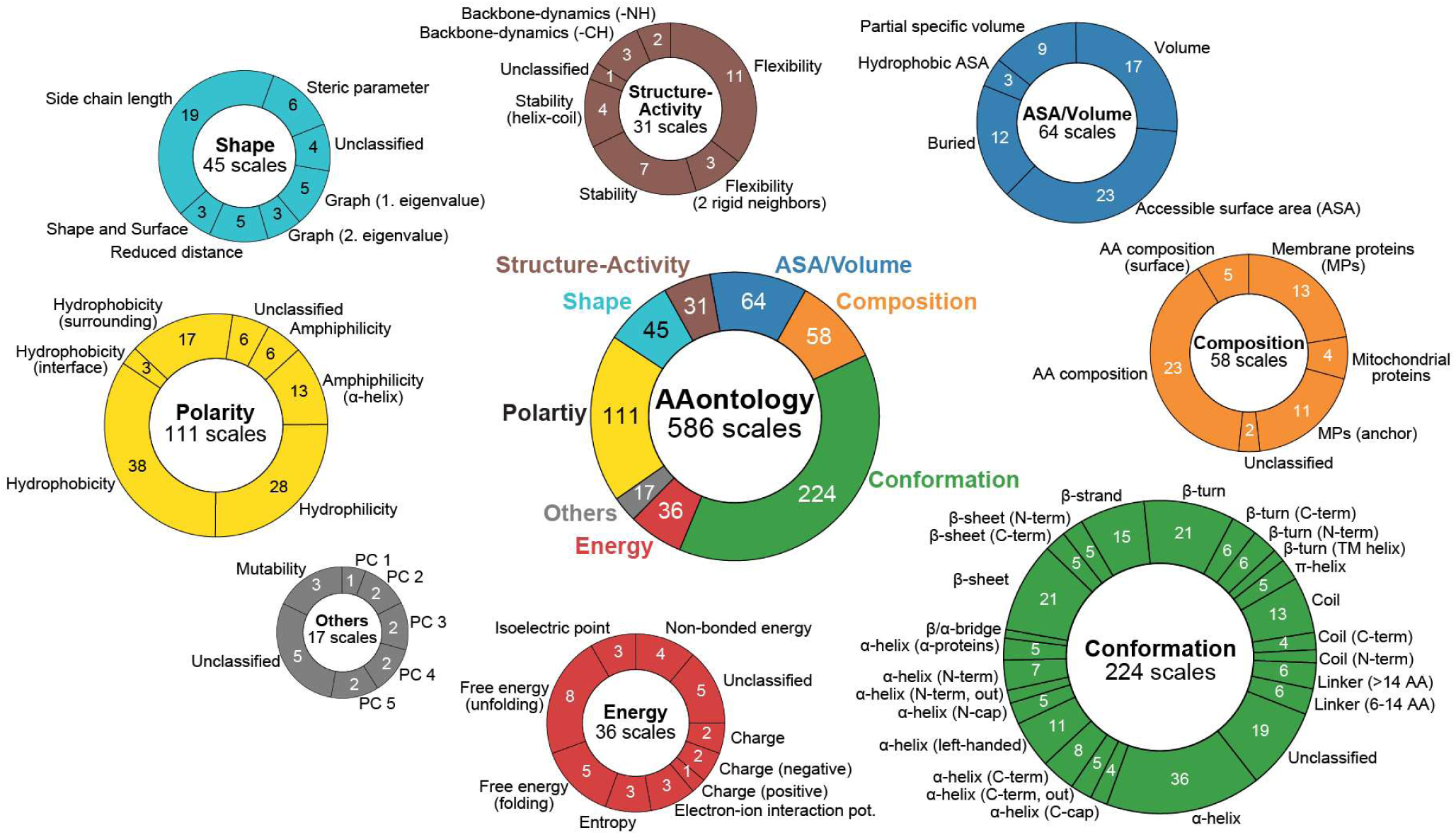
AAontology: Categories and subcategories. Two-level hierarchy of AAontology comprising 67 subcategories within the following 8 categories (excluding unclassified scales): ‘ASA/Volume’ (blue; ‘ASA’: Accessible Surface Area) with 5 subcategories, ‘Composition’ (orange) with 5 subcategories, ‘Conformation’ (green) with 24 subcategories, ‘Energy’ (red) with 9 subcategories, ‘Others’ (gray) with 6 subcategories, ‘Polarity’ (yellow) with 6 subcategories, ‘Shape’ (light blue) with 6 subcategories, and ‘Structure-Activity’ (brown) with 6 subcategories. Each category is visualized by a donut plot, with its size approximately reflecting the number of scales assigned to the respective category.

### 3.1 Scale categories

Scale subcategories were manually named using our bag-of-word analysis as guidance and consolidated based on selected studies. Brief descriptions of each scale, scale subcategory, and scale category can be found in **Supplementary Table 3**. Each category (**Fig. 5–Fig. 12**) including their subordinated scale subcategories will be described in the following.

**Fig. 5.**
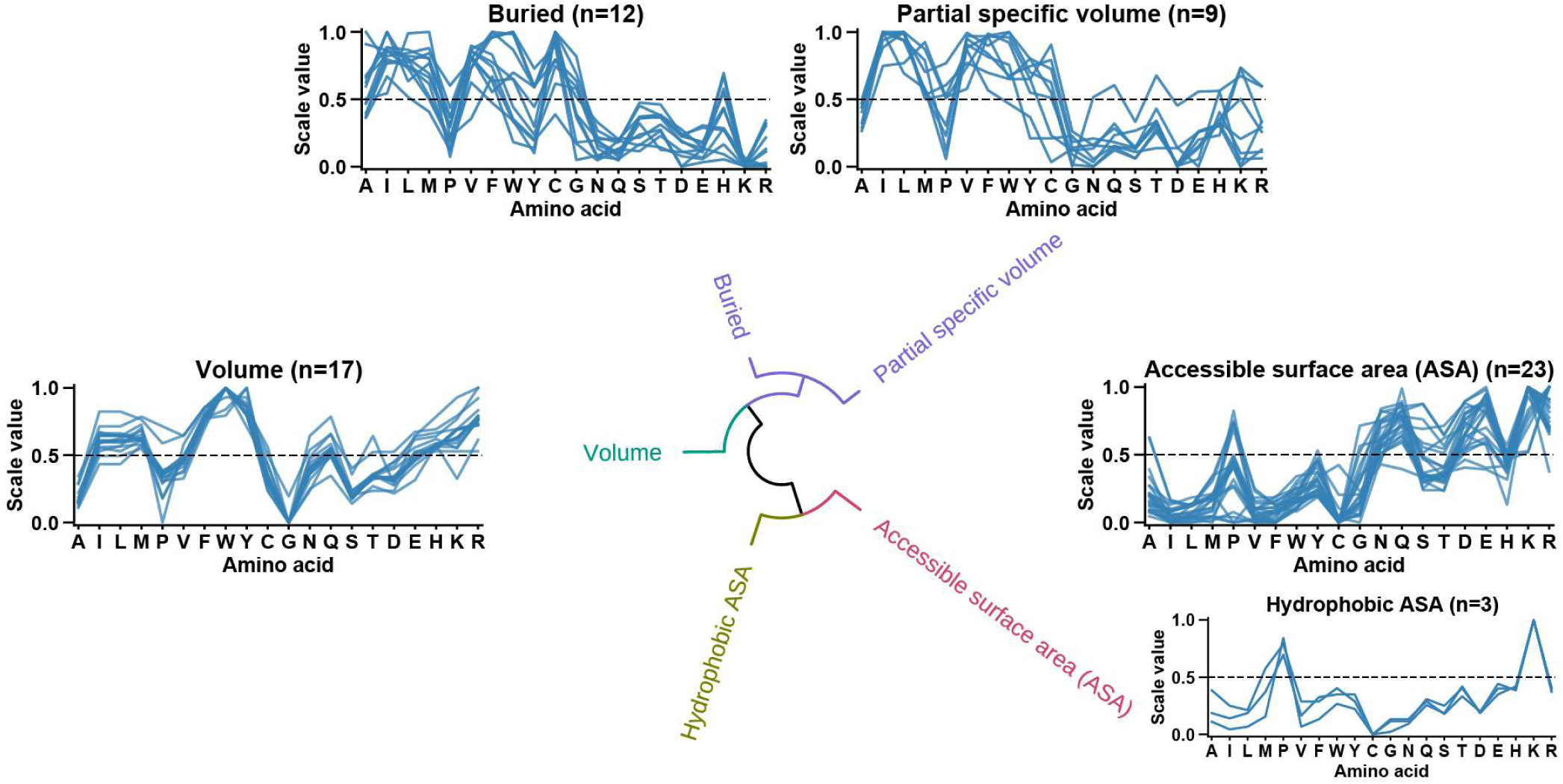
ASA/Volume category. Circular dendrogram showing the hierarchical clustering of all ‘ASA/Volume’ subcategories, based on the Euclidean distance between their average scales. Each subcategory is represented by a line plot, with scales depicted as lines and the number of scales assigned to a subcategory indicated in parentheses. The arrangement of line plots corresponds to the clustering results, and size modifications have been made if necessary, highlighting larger subcategories. Amino acids are given by their one-letter code and are grouped as follows: non-polar/hydrophobic amino acids comprising alanine (A), isoleucine (I), leucine (L), methionine (M), proline (P), and valine (V); aromatic amino acids of phenylalanine (F), tryptophan (W), and tyrosine (Y); polar/hydrophilic amino acids comprising cysteine (C), glycine (G), asparagine (N), glutamine (Q), serine (S), and threonine (T); acidic/negatively charged amino acids of aspartic acid (D) and glutamic acid (E); as well as basic/positively charged amino acids of histidine (H), lysine (K) and arginine (R).

**Fig. 6.**
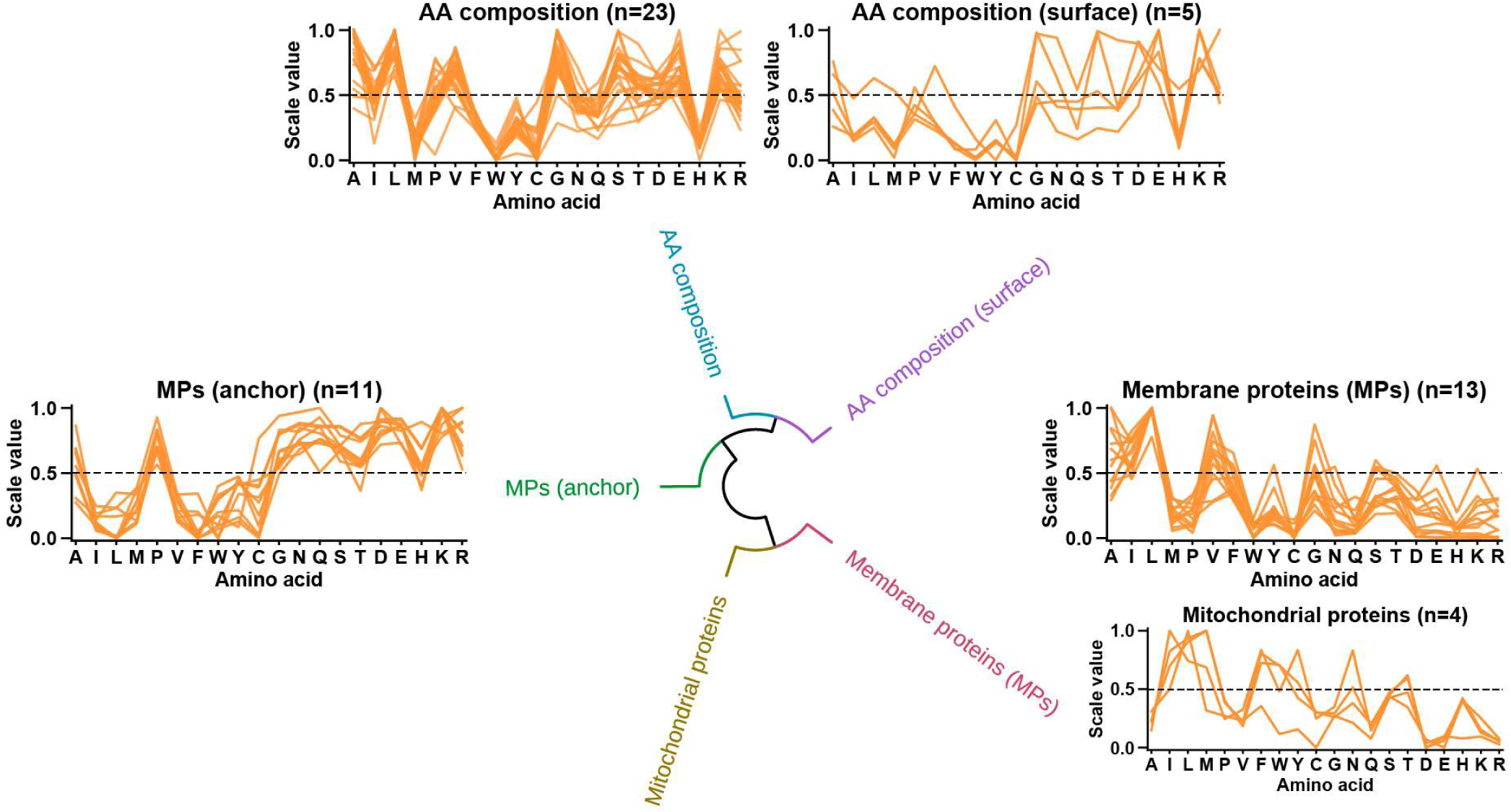
Composition category. Circular dendrogram showing hierarchical clustering of all ‘Composition’ subcategories. Further details are as described in **Fig. 5**.

**Fig. 7.**
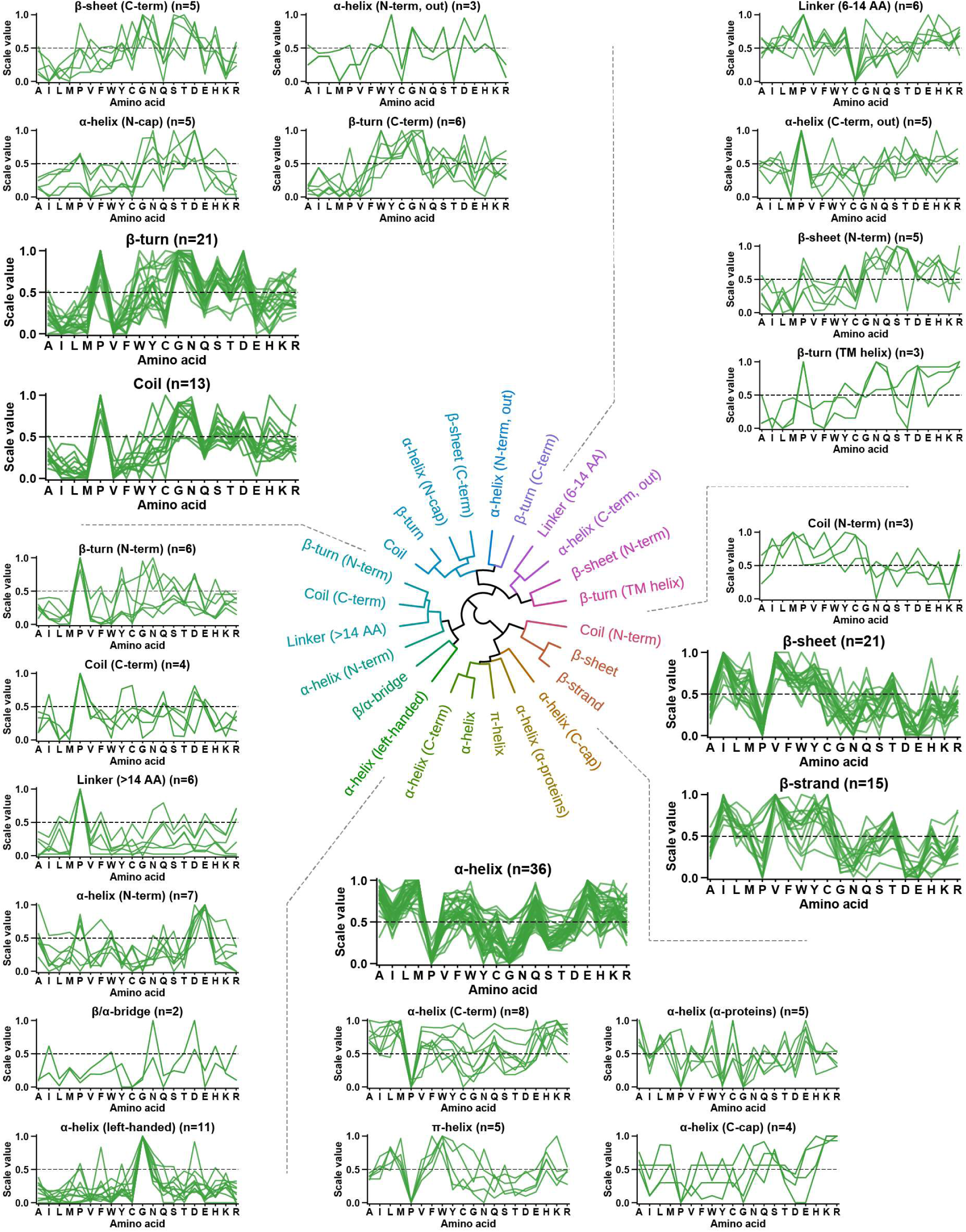
Conformation category. Circular dendrogram showing hierarchical clustering of all ‘Conformation’ subcategories. Grouping of line plots representing subcategories follows clustering results, as highlighted by dashed gray lines. An explanation of further details is given in **Fig. 5**.

**Fig. 8.**
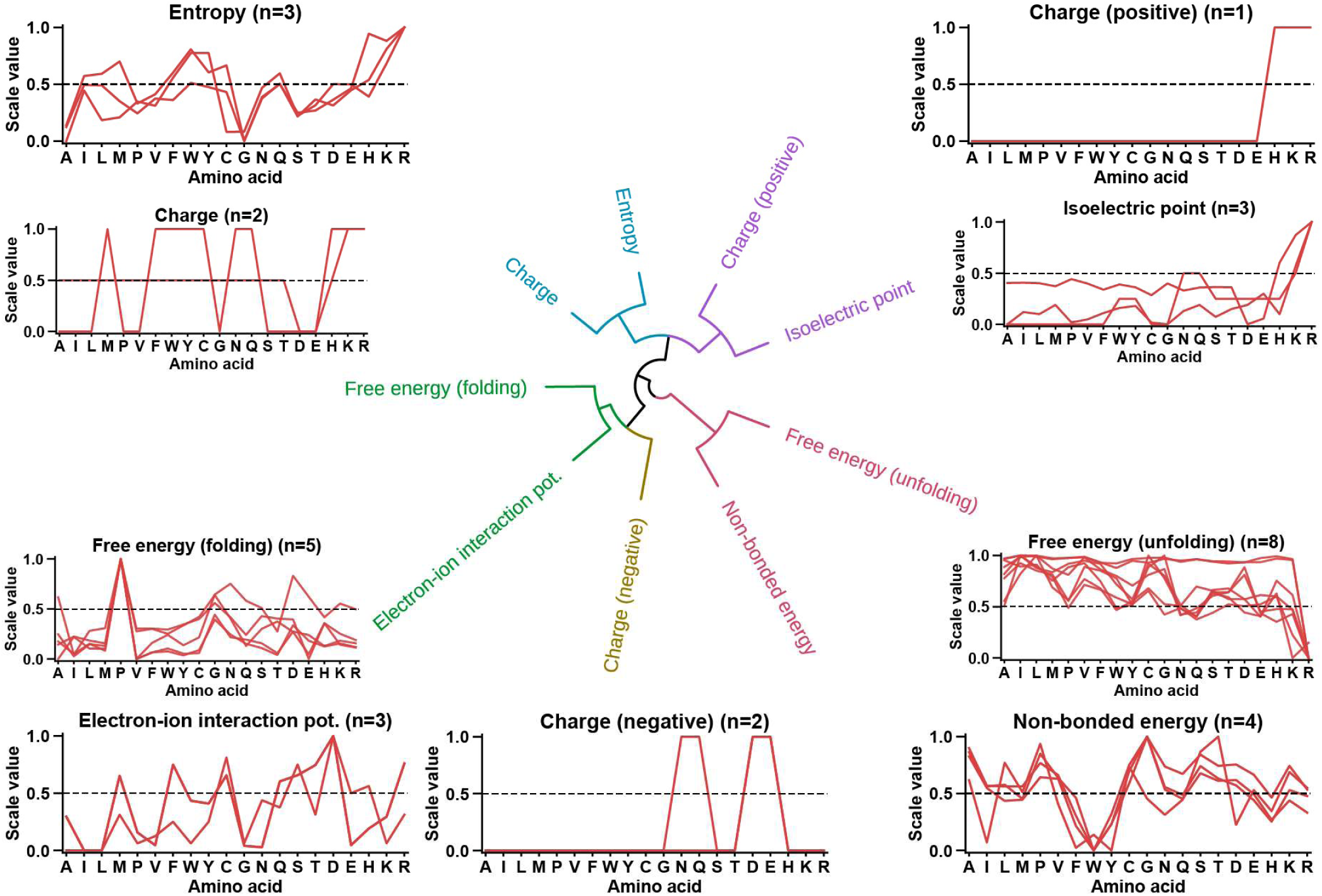
Energy category. Circular dendrogram showing hierarchical clustering of all ‘Energy’ subcategories. Further details are given in **Fig. 5**. Note that we corrected the following transcription mistake in AAindex database for the ‘charge transfer capability’ [123] scale (CHAM830107 scale id; *Charge (negative)* subcategory): glycine was erroneously scored 1 instead of glutamine.

**Fig. 9.**
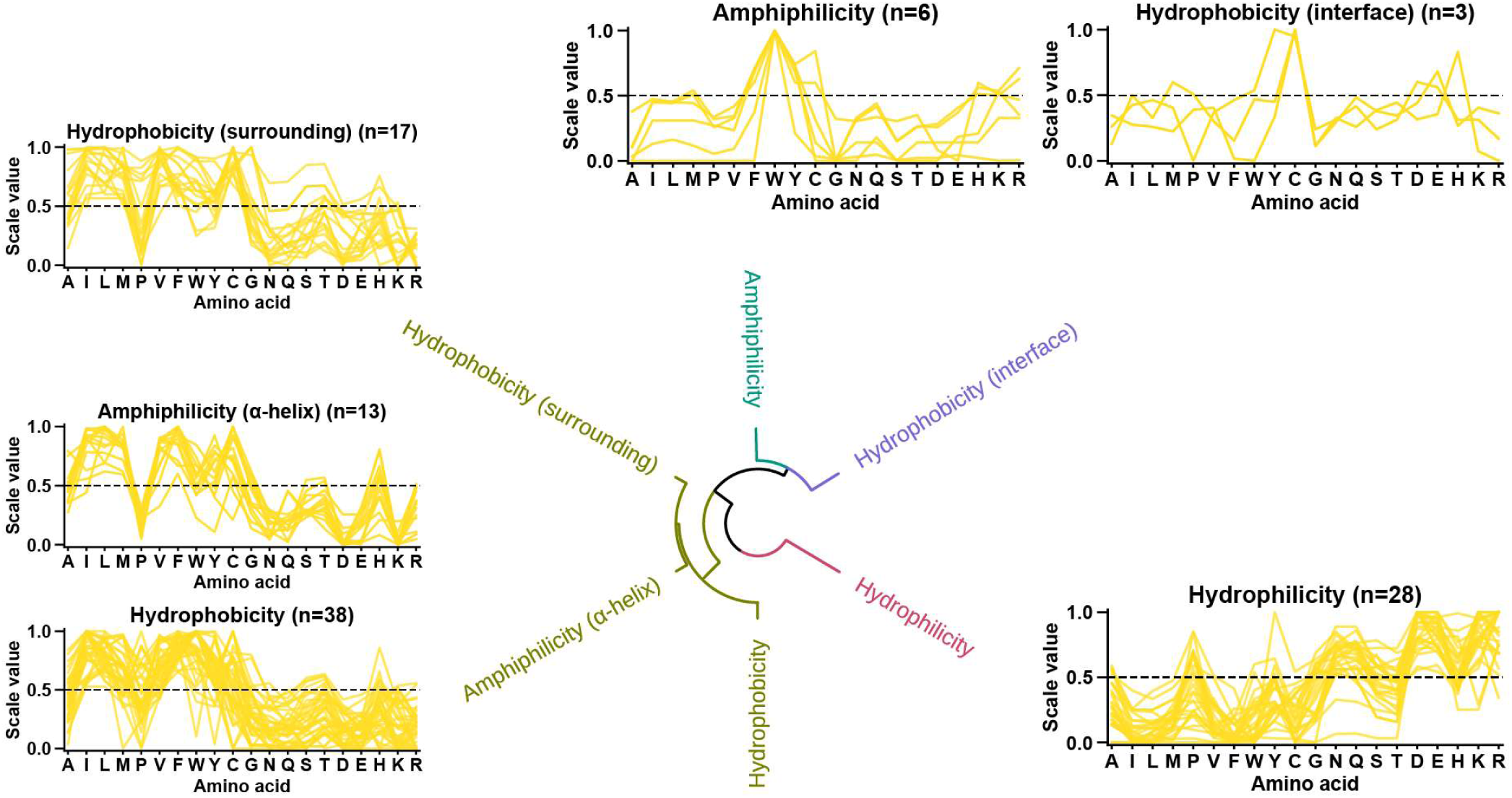
Polarity category. Circular dendrogram showing hierarchical clustering of all ‘Polarity’ subcategories. See **Fig. 5** for further details.

**Fig. 10.**
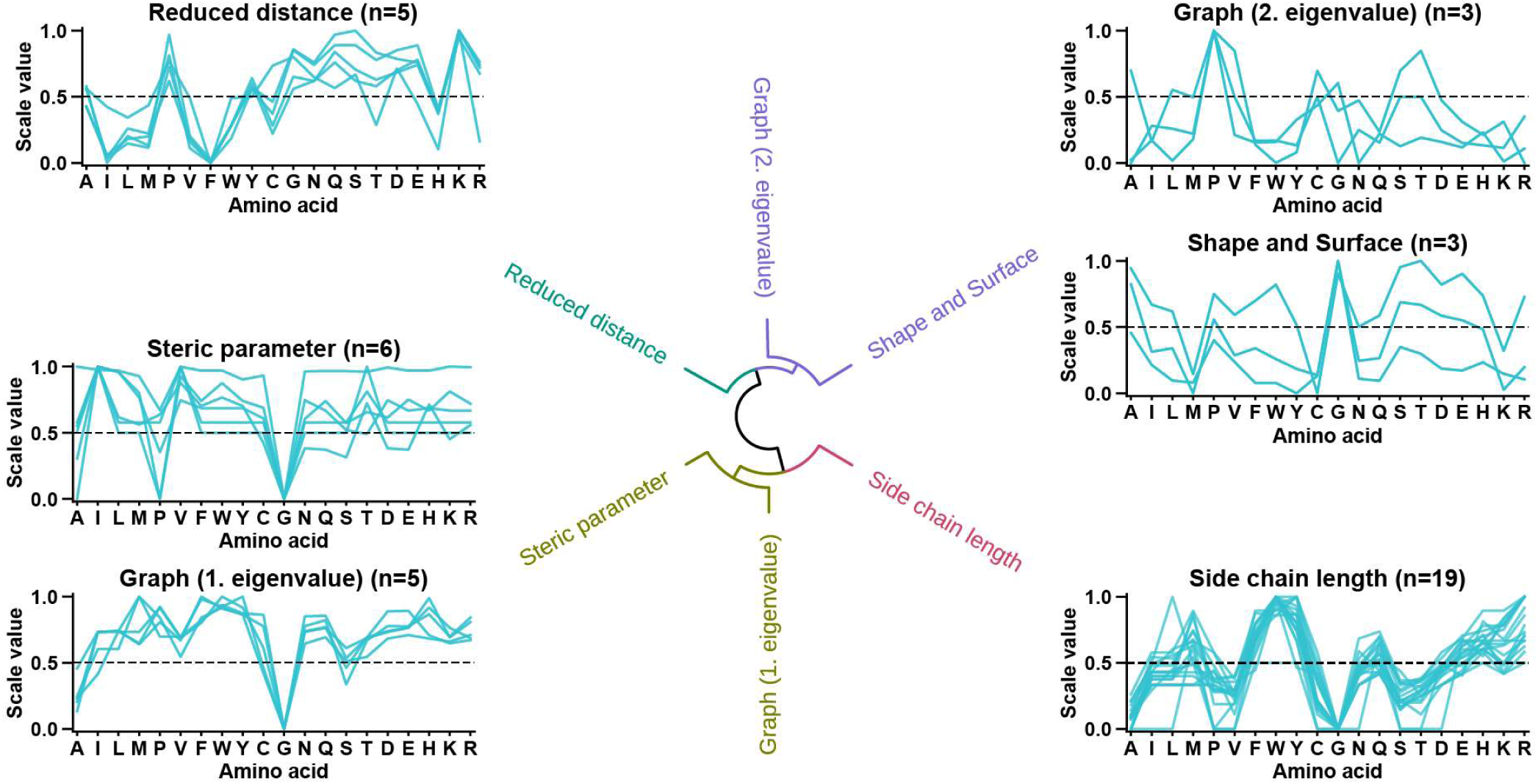
Shape category. Circular dendrogram showing hierarchical clustering of all ‘Shape’ subcategories. More details are explained in **Fig. 5**.

**Fig. 11.**
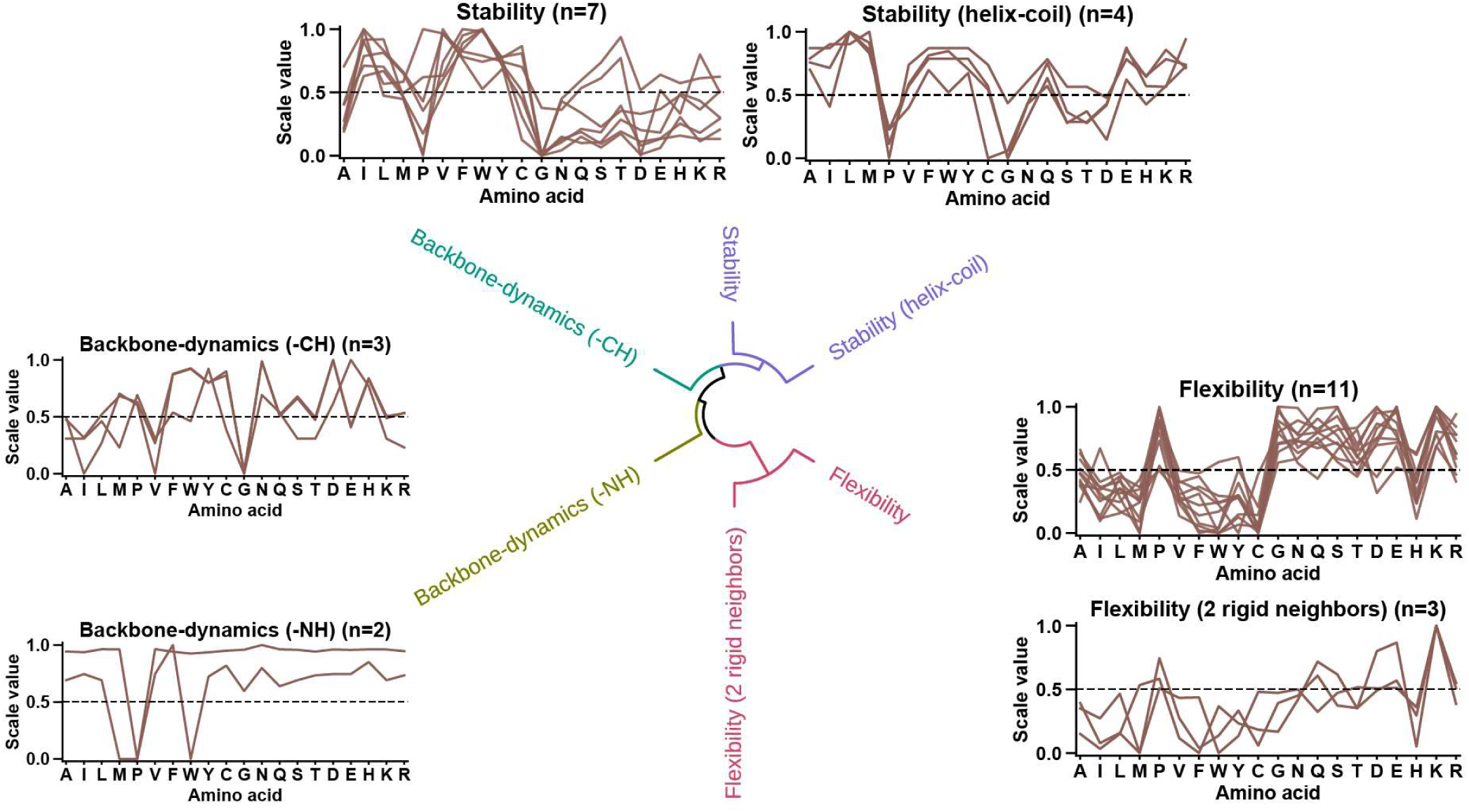
Structure-Activity category. Circular dendrogram showing hierarchical clustering of all ‘Structure-Activity’ subcategories. Details are described in **Fig. 5**.

#### 3.1.1 Accessible surface area (ASA)//Volume

The ‘ASA/Volume’ category comprises around 60 scales and 5 subcategories describing properties related to the general volume of amino acids and their preference of being either accessible to solvent, reflected by their ASA, or being not accessible to solvent, *i.e.*, being buried within a folded protein.

*Accessible Surface Area (ASA)* (n=23) measures the residue surface area that is accessible/exposed to solvent (typically water), obtained from folded proteins. It indicates the ability of residues to interact with water, mainly at the protein surface. Residues with larger ASA often participate in protein-protein interactions, with higher ASA being more typical for polar residues with longer side chain (lysine > arginine > glutamic acid > glutamine) [44,54].

*Hydrophobic ASA* (n=3) measures the residue surface area that is solvent-accessible and hydrophobic, obtained from folded proteins. This reflects the hydrophobic area exposed to water at the protein surface. This value is remarkably high for lysine—due to its long hydrophobic side chain ending in a polar ε-amino group—followed by proline, a hydrophobic residue often found in β-turns on the protein surface. Conversely, aromatic residues show moderately low values due to their less frequent occurrence on the protein surface [44].

*Buried* (n=12), as opposed to ASA, represents the propensity of amino acids to be buried within the protein core, shielded from the exterior solvents. These concealed residues are crucial for the protein stability and folding, driven by hydrophobic interactions and disulfide bridges. Consequently, cysteine and large hydrophobic residues tend to be particularly buried [54,55].

*Volume* (n=17) is a direct measure of the amino acid size. Larger amino acids can enhance protein-protein interactions [56] and foster protein stability through long-range interactions [57]. However, their size requirements can impact chain packing [58]. This becomes crucial when mutations alter amino acid size in densely packed protein cores, which can lead to destabilization [59]. Aromatic amino acids (tryptophan > tyrosine > phenylalanine) and arginine are the largest, while glycine and alanine are the smallest [60].

*Partial specific volume* (n=9) reflects the effective amino acid volume in water , accounting for both physical volume and additional water displacement due to residue-solvent interactions [61]. Also included in this subcategory are hydrophobic interactivity potential [62] and bulkiness (*i.e.*, the side chain volume/length ratio) [63]. These properties affect protein structure by promoting stability and introducing steric hindrances. Arginine, despite its high *Volume*, exhibits only a moderate *Partial specific volume* as it lacks additional hydrophobic water displacement. In contrast, large hydrophobic residues, particularly branched-chain amino acids (isoleucine, leucine, valine) [64] and aromatic residues (phenylalanine, tryptophan, tyrosine), score highest [62,63,65].

#### 3.2.2 Composition

The ‘Composition’ category includes around 60 scales and 5 scale subcategories regarding the frequency of amino acid occurrence in different types of proteins, such as membrane proteins or the mitochondrial proteins.

*AA Composition* (n=23) represents the overall frequency of amino acids (abbreviated here as ‘*AA*’) in proteins, largely independent of subcellular location or the residue position within a protein. This subcategory also comprises specific compositions, such as those of intra- or extracellular proteins, that correlate strongly with the general composition. Most abundant are alanine, leucine, and glycine, while amino acids containing a sulfur atom (methionine, cysteine) or a nitrogen atom within an aromatic ring (tryptophan, histidine) occur less frequently [66–68].

*AA Composition (Surface)* (n=5) reflects the propensity of amino acids to occure at the protein surface compared to the protein interior. Polar amino acids typically occur frequently on the protein surface, with aspartic acid and lysine being the most prevalent [69].

*Membrane Proteins (MPs)* (n=13) describes the amino acid frequency in transmembrane domains. These α-helical domains traverse cellular membranes, forming parts of either single-spanning or multi-spanning membrane proteins. Given their specialized roles within lipid-rich environments, these domains exhibit a distinct composition, predominantly non-polar amino acids (leucine > valine > alanine, phenylalanine), with methionine and tryptophan appearing least frequently [70,71].

*MPs (anchor)* (n=11) refers to the frequency of amino acids occurring in the N-/C-terminal regions flanking the transmembrane domains (TMD) of membrane proteins. The flanking regions fortify the anchoring of the hydrophobic TMDs [72] through strong partitioning between the hydrophobic and hydrophilic phase [73,74]. They terminate TMD helices by short motifs of hydrophilic amino acids, typically comprising charged helix-capping and/or helix-breaking residues (proline and glycine) [75]. The N-terminal region is characterized by negatively charged/acidic residues, especially aspartic acid but also asparagine. In contrast, the C-terminal region follows the positive-inside rule [76], with poly-basic motifs (arginine, lysine) enhancing protein-phospholipid interactions at the interface [73,75,77].

*Mitochondrial Proteins* (n=4) focuses on the frequency of occurrence of amino acids in mitochondrial proteins. Similar to membrane proteins, they are rich in hydrophobic residues, but typically with less valine [70].

#### 3.2.3 Conformation

The ‘Conformation’ category is the largest category, with over 200 scales across 24 subcategories. It covers four major conformations: helical, extended (β-sheet and β-strand) together with β-turn, and coil secondary structures. These account for almost all secondary structures—roughly 30–40% α-helix, 20– 30% β-sheets, 20% β-turn, and 20% coils [78]. Pioneering work by Chou and Fasman [79,80], Richardson and Richardson [81,82], as well as Qian and Sejnowski [83] highlighted the distinct amino acid distribution across and within these conformations.

Helical and extended conformations are structured by hydrogen-bonding patterns, adopting specific dihedral angles within the polypeptide backbone [84]. Conversely, coils are defined by the absence of these patterns according to the Define Secondary Structure of Proteins (DSSP) convention [85]. Coils can either adopt disordered conformations (called ‘random’ coils) or form structured loops (non-random coils [86]) in folded proteins. Secondary structure formation is context-dependent, demonstrated by conformational transition studies for polyalanine [87] and polylysine [88]. With decreasing temperature, the conformational prevalence order is described by coil > β-sheet > helix, whereas an decrease in solvent hydrophobicity reorders this preference to coil > α-helix > β-sheet [87,89]. For example, in hydrophilic solvents, α-helices are less stable than β-sheets but vice versa in hydrophobic membranes. α-Helices can segregate hydrophobic and hydrophilic amino acids, while β-sheets shield hydrophobic residues from water [44], albeit playing a multifaceted role in protein aggregation [90]. Intriguingly, proline and glycine (to a lesser extent), abundant in random coils, can interrupt α-helix formation and increase flexibility [91]. In β-sheets, these residues can destabilize the central structure but enhance peptide chain turns at the edges.

Each conformational subcategory describes the prevalence/tendency of residues to occur within the associated secondary structure. However, for the sake of clarity, only the secondary structure and their conformational implications will be described, while bearing in mind that these subcategories essentially represent residue frequencies.

##### Helical conformations

These subcategories describe α-helical and π-helical conformations, differing in the number of residues per 360° turn—3.6 and 4.4, respectively. The more compact 310-helix, comprising 3.0 residues per turn, is not described in AAindex, and thus not included here. Although 310-helices are rare (4%) due to their reduced stability [92,93], short 310-helix segments often appear at the C-terminus of α-helices, tightening the final helical turn [81]. They can also switch entirely to an α-helical conformation [92]. The compactness order of these helix types is 310-helix > α-helix > π-helix, each characterized by distinct backbone hydrogen bonding patterns of *i* to *i*+3, *i*+4, *i*+5, respectively [94]. They exhibit an altered stability order of α-helix > 310-helix > π-helix, explaining their respective prevalences of 30-40%, 4%, and 0.02% [92]. The π-helix prevalence might, however, be higher than suspected [93], particularly for short (7 to 10 residue long) π-helical segments [95].

*α-helix* (n=36) refers to right-handed helical structures with 3.6 residues per turn. It is the most abundant protein helix type, accounting for 30–40% of all secondary structures [93]. α-Helices are formed by backbone hydrogen bonding (*i*+4) and local side chain interaction between periodically neighbouring residues (*i*+3/4), which increase helical stability via hydrophobic, polar, and aromatic stacking interactions [96–98]. Predominant residues include alanine, large hydrophobic residues (leucine, methionine), and glutamic acid alongside glutamine [80].

*α-helix (N-term)* and *α-helix (C-term)* (n=7, 8) denote the segments at the N-terminus and C-terminus inside the α-helix. These segments, especially the terminal residues, can notably influence the stability of the α-helix structure [75,80,82].

*α-helix (N-cap)* and *α-helix (C-cap)* (n=5, 4) refer to the positions at the exact termini of the α-helix where the helix is ‘capped’, *i.e.*, either begins (N-cap) or ends (C-cap), respectively. The N-terminus has a high prevalence for aspartic acid, but also asparagine and serine, while the C-terminus is characterized by positively charged/basic residues (arginine, lysine). These residues link α-helices to adjacent turns or unstructured regions [75,82,99].

*α-helix (N-term, out)* and α-helix *(C-term, out)* (n=3, 5) refer to the segments at the N-terminus and C-terminus outside the α-helix structure, critical for the termination of the helix. The N-terminus often harbors negatively charged/acidic residues (mainly aspartic acid) but also residues with large side chains (mainly tyrosine) and the helix-breaking glycine. In contrast, the C-terminus comprises positively charged/basic residues (histidine, arginine), residues with a large side chain (mainly phenylalanine), and, especially, the helix-breaking proline [75,82].

*α-helix (α-proteins)* (n=5) reflects the frequency of residues in α-helices within proteins that predominantly have α-helices as their secondary structural element [100].

*α-helix (left-handed)* (n=11) is a left-handed helical structure with 3.6 residues per turn, similar to right-handed helices, but winds in the opposite direction [84], resulting in steric hindrance between side chain and main chain atoms. Although rare and typically 4-residue long, these helices contribute to protein stability and functions such as ligand binding or active site participation [101]. Glycine and aspartic acid are prevalent [102].

*π-helix* (n=5) is a rare and unstable [92] helix with 4.4 residues per turn, resulting in a 1 Å hole at its center too narrow to accommodate a water molecule. This hole diminishes backbone van der Waals interactions, destabilizing π-helices [94]. Mainly stabilized by side-chain interactions, including aromatic stacking and van der Waals forces, π-helices require a collinear alignment of every fourth side chain residue [94]. This unique structure, found in specific protein binding sites [94], may enhance enzyme ligand coordination and dipole involvement in ion transport proteins [93]. Prevalent residues include aromatic and large aliphatic amino acids—particularly tryptophan, phenylalanine, and methionine [93].

##### Extended conformations and β-turn

These subcategories comprise extended conformations (*i.e.*, β-strand and β-sheet) and β-turn, a sharp 4 residue turn often linking consecutive segments within anti-parallel β-sheets, which are characterized by adjacent β-strands running in opposite directions.

*β-strand* (n=15) refers to a fully extended segment of a polypeptide chain, typically containing 3 to 10 residues. When multiple β-strands align, they form β-sheets, which are predominantly intra-molecular. Inter-molecular β-sheets (‘hybrid β-sheets’ [103,104]) can also mediate protein-protein interactions, facilitated by residue side chains fostering interaction specificity and affinity [105]. Prevalent residues are branched-chain amino acids (value, isoleucine > leucine) and cysteine [80,106]. *β-sheet* (n=21) is, after *α-helix*, the second most common secondary structure, characterized by its pleated structure, also called pleated sheet. It is composed of multiple β-strands forming distinct backbone hydrogen bonding patterns (either anti-parallel or parallel) [107], supported by hydrophobic side chain interactions and avoidance of steric side/main chain clashes [108]. β-Strands are connected by short loops of two to five residues or β-turns. Prevalent residues are branched-chain (valine, isoleucine > leucine) and aromatic (phenylalanine, tyrosine > tryptophan) amino acids [80,83].

*β-sheet (N-term)* and *β-sheet (C-term)* (n=5, 5) denote the N-terminal and C-terminal segment of a β-sheet, respectively. Generally, prevalent residues near or at the termini are serine and the destabilizing proline (inducing a 90° backbone turn) and glycine (enhancing backbone flexibility) [108]. The C-terminus additionally contains asparagine and aspartic acid, while the N-terminus is also characterized by positively charged residues (histidine, arginine) [83].

*β/α-bridge* (n=2), a new term introduced here, pertains to the frequency of residues in the ‘bridge region’ of the Ramachandran plot [84], an area reflecting an energetically less favored conformation between β-sheet (top-left quadrant) and right-handed α-helix (bottom-left quadrant). Higher scores suggest that these residues can facilitate context-dependent folding, such as α-helix/β-sheet transition, by acting as a conformational switch by engaging in long-distance (intra)molecular hydrogen bonding [109–111]. Polar residues with longer side chains and moderate β-sheet or α-helix preferences are favored, *e.g.*, asparagine, aspartic acid, and histidine [102,112,113].

*β-turn* (n=21), also known as β-bend or hairpin loop, is the third most common secondary structure. Comprising 4 amino acids, β-turns sharply reverse the polypeptide chain by 180°. This reversal is often aided by a hydrogen bond between the 1st and the 4th residue, yielding multiple subclassifications [114]. Typically connecting consecutive segments within anti-parallel β-sheets, β-turns can overlap with other structural elements (*e.g.*, co-occurrence of 1st residue in α-helix or β-sheet [78]) and frequently appear on protein surfaces, contributing to compact structures. Prevalent residues include proline, glycine, asparagine, and aspartic acid [80,115].

*β-turn (N-term)* and *β-turn (C-term)* (n=6, 6) respectively represent the prevalence of residues for the 1st and 2nd positions, and the 3rd and 4th positions within the β-turn structure [80,102].

*β-turn (TM helix)* (n=3) refers to β-turns placed in the middle of a long single-spanning transmembrane helix. Unlike α-helical turns, completing 360° over 3 to 4 residues, β-turns accomplish a 180° turn over 4 residues. This chain reversal can transform a single long transmembrane helix into two closely spaced helices. Prevalent are charged residues (aspartic acid, arginine > glutamic acid, lysine) and the helix-breaking proline, while glycine, compared to its presence in *β-turn*, demonstrates only a moderate propensity [116].

##### Coil conformation and linkers

These subcategories describe coil conformations and linker segments. Unlike helical and extended secondary structures, coils, devoid of a defined backbone hydrogen-bonding pattern, can adopt either unstructured coil (‘random coils’) or structured loop forms. Linkers, typically unstructured in isolation, may assume specific conformations upon interaction with other protein parts or under certain conditions. While random coils and unstructured linkers lack a defined structure, their roles in proteins differ. Coils are highly dynamic and can mediate various functions, whereas linkers specifically connect functional domains, providing flexibility.

*Coil* (n=13), the fourth major secondary structure, is defined by the absence of well-defined backbone hydrogen-bonding patterns [84], which are characteristic for α-helices or β-sheets. Coils can be ‘random’ coils, typically elusive in X-ray crystallography, or structured loops (non-random coils [86]), which are observable [117]. ‘Random’ coils are dynamic, flexible segments enabling critical conformational changes in proteins and playing a vital role in molecular interactions, especially in intrinsically disordered proteins [118,119]. In contrast, structured loops connect α-helix or β-sheet motifs in folded proteins [120], contributing to molecular recognition at the protein surface [117]. Akin to β-turns [121], prevalent residues are proline, glycine, asparagine, and serine [83,115].

*Coil (N-term)* and *Coil (C-term)* (n=3, 4) denotes the N-terminal and C-terminal sections of coil conformations, respectively. Notably, methionine is typically present in the N-terminus but absent at the N-terminus of a coil [83].

*Linker (>14 AA)* and *Linker (6-14 AA)* (n=6, 6) describes regions that connect functional domains, called linkers. The linker length significantly influences protein structure and function, enabling cooperative interactions between domains. Long linkers (>14 residues) predominantly adopt helical or coil structures, ensuring flexibility and domain separation. Medium-sized linkers (6–14 residues), however, can also form β-strand structures, striking a balance between flexibility and stability [122].

#### 3.2.4 Energy

The ‘Energy’ category comprises around 40 scales organized into 9 specific subcategories, each highlighting different energetic aspects of amino acids including free energy—determining conformational stability—and charge, playing an important role for protein structure and function such as enzymatic activity, protein interactions, or anchoring of transmembrane proteins.

*Charge* (n=2) represents net charge and ’charge transfer donor capability’ [123]. The net charge reflects proton-related ionic charge, assigning scores 1, 0, 0.5 to positively charged (arginine, lysine), negatively charged (aspartic and glutamic acid), and all other residues, respectively. [124]. The ‘Charge transfer donor capability’ marks the ability (1) or inability (0) to donate an electron with a certain ionization energy input. It is present in polar (cysteine, asparagine, glutamine), basic (histidine, arginine, lysine), and aromatic residues, as well as in methionine (due to its sulfur atom) [123].

*Charge (negative)* (n=2) describes a residués negative charge (aspartic acid, glutamic acid) [125] and ‘charge transfer capability’ [123]. The latter involves polar side chain groups, specifically CONH2 in asparagine and glutamine and the COO- in deprotonated aspartic and glutamic acids, crucial for intermolecular interactions such as hydrogen bonding [126]. While the deprotonated aspartic and glutamic acid can accept protons [125], asparagine and glutamine can accept electrons [123,125]. Presence is indicated as 1, absence as 0 [123,125].

*Charge (positive)* (n=1) is dedicated to residues carrying a positive charge (arginine, lysine, histidine), indicating presence with 1 and absence with 0 [125].

*Entropy* (n=3) refers to side chain conformational entropy, reflecting the number of potential conformations a residue can be part of. It is one opposing force to protein folding [127,128] and it also influences molecular interactions [129]. Alanine has a low *Entropy* due to its predominant α-helix propensity, while structure-breaking residues (glycine > proline) also exhibit low *Entropy*. In contrast, positively charged and aromatic residues have high *Entropy* due to their moderate propensities for various secondary structures such as α-helices or β-sheets [130].

*Free energy (unfolding)* (n=8) encompasses measures of conformational stability, including the ‘Gibbs energy of unfolding’ and ‘activation Gibbs energy of unfolding’ [131]. Higher values indicate enhanced stability and a larger energy barrier against unfolding. The energy required for residue transfer from hydrophobic to hydrophilic solvent, another component, reflects role of hydrophobicity role in stability [132]. Despite consistent high activation energy across residues, excluding arginine, other measures are more nuanced, with positively charged residues (arginine > lysine > histidine) scoring higher [131,132].

*Free energy (folding)* (n=5) comprises measures of the absolute free energy required for α-helix or β-strand formation, where higher values denote reduced stability, hence conformational instability [133]. While structure-disrupting residues (proline > glycine) show high values, asparagine and aspartic acid display a stronger tendency to destabilize β-strands [134].

*Isoelectric point* (n=3), often abbreviated as pI, designates the pH at which an amino acid is electrically neutral, indicating relative acidity or basicity. Basic residues (arginine > lysine > histidine) exhibit the highest values, while acidic residues show the lowest [63]. Additionally, residue basicity is also determined based on hydrogen bond donation, crucial for enzymatic reactions and protein-ligand interactions [135]. Quantified as the number of hydrogen bond donors within a side chain [125], this parameter ranges from none (non-polar residues) to four (arginine), where each donated bond equates to 0.25 [63,125].

*Electron-ion interaction potential* (n=3) denotes the capacity for electrostatic interactions, such as dipole-dipole, hydrogen bond, or ionic interactions. It is quantified by the average energy state of all valence electrons, the electrons within the outer shell of atoms. Essentially, it measures the potential of amino acids for engaging in electrostatic protein-protein and protein-DNA interactions [136]. Aspartic acid exhibits the highest score, followed by cysteine and serine, while glycine and the branched chain amino acids show the lowest scores [137].

*Non-bonded energy* (n=4) refers to the average energy per residue resulting from non-covalent interactions. Derived from X-ray crystallography using the Lennard-Jones potential, this energy encompasses electrostatic interactions and van der Waals forces, with the latter not being considered in the *Electron-ion interaction potential*. Therefore, smaller residues (glycine > proline > alanine) score highest due to their ability for close packing, fostering van der Waals interactions. Conversely, larger aromatic amino acids yield lower scores, as their size dilutes the energy contribution per atom [138].

#### 3.2.5 Polarity

The ‘Polarity’ category is the second largest category with over 100 scales organized into 6 subcategories. This category describes the foundational dichotomy of hydrophilicity and hydrophobicity, reflected by polar and non-polar residues which determine crucial biological phenomena such as protein folding and subcellular localization.

*Hydrophilicity* (n=28) reflects the preference of an amino acid for a polar/hydrophilic environment. This is often gauged as the transfer free energy from water to a non-polar solvent. It is essential for protein-solvent interactions, influencing protein solubility, folding, and function. Charged residues exhibit the highest hydrophilicity [132,139].

*Hydrophobicity* (n=38) represents the amino acid preference for a non-polar/hydrophobic environment. It is often measured as the transfer free energy from a non-polar solvent to water or from the interior of a protein to the surface, playing a central role in protein folding and stability. Isoleucine and phenylalanine consistently score highest across all scales [132,139,140].

*Hydrophobicity (surrounding)* (n=17) describes the effective hydrophobicity of residues within globular proteins, accounting for both their own hydrophobicity and the hydrophobicity of the neighbouring residues within an 8-angstrom radius. Derived from protein crystal structures, it reflects *Hydrophobicity* in internal protein arrangements adjusted for steric hindrance, thereby gauging protein stability. Compared to *Hydrophobicity*, it is closer related to *Buried* measures, with particularly high scores for cysteine, slightly lower ones for aromatic residues, and similarly high scores for branched-chain amino acids (valine > isoleucine, leucine) [141].

*Hydrophobicity (interface)* (n=3) focuses on the preference of residues for non-polar/hydrophobic environments at membrane interfaces, influencing the membrane-association behavior of proteins [142]. Cysteine exhibits the highest scores, followed by tyrosine. [45].

*Amphiphilicity* (n=6) captures the preference of amino acids to occur at the interface of polar and non-polar solvents, such as the membrane-water boundary. Typically, high at the termini of transmembrane helices, this can substantially affect protein interactions and cellular processes. Amphiphilic residues consist generally of a polar and a non-polar group (alkyl side chain or aromatic ring). Tryptophan scores notably higher than other residues, followed by tyrosine and basic amino acids (arginine > lysine, histidine) [143].

*Amphiphilicity (α-helix)* (n=13) denotes the amino acid propensity to form amphiphilic α-helices, characterized by segregated polar and non-polar faces [144,145]. These helices, when located at protein surfaces or membranes, often serve as signal sequences [146]. When interacting with membranes, they align parallel to the membrane, discerning curvature and aiding remodeling [147]. Prevalent are non-polar residues (methionine, tryptophan, and branched chain amino acids), along with certain polar (cysteine > tyrosine) and positively charged residues (histidine > arginine) [146,148].

#### 3.2.6 Shape

The ‘Shape’ category, embracing 45 scales across 6 subcategories, delves into geometric and steric characteristics of amino acids. These subcategories describe side chain angles, symmetry, and unique parameters derived from amino acid representation based on graph theory. Here, amino acids are conceptualized as undirected, node-weighted graphs, with atoms as nodes and molecular bonds as edges [149]. From these graphs, different measures can be derived, such as the maximum eccentricity—the greatest number of bonds (edges) required to link the two furthest atoms within a graph. For instance, this results in 0 for glycine, 1 for alanine, or 3 for serine or cysteine. These metrics offer a mathematical fingerprint of each amino acid’s atomic structure.

*Side chain length* (n=19) refers to the length of the amino acid side chain, quantified by the number of bonds in its longest chain. This subcategory includes also closely related graph-based size measures such as the average eccentricity, offering an alternative perspective on the amino acid length [123,149]. *Graph (1. eigenvalue)* and *Graph (2. eigenvalue)* (n=5, 3) are graph-based measures, where amino acids are considered as undirected graphs, with atoms as nodes and bonds as edges. To capture the edge-bond relationships, the Laplacian matrix is computed from these graphs. Of interest are this matrix’s first (*i.e.*, smallest) and second smallest eigenvalues. These measures allow a nuanced differentiation between amino acids, even those with similar atomic configurations [149].

*Reduced distance* (n=5) reflects the relative distance of a residue from the protein’s center of mass, as obtained for 14 native proteins [150]. To adjust for protein size and shape, this distance is divided by the root mean square of the ‘radius of gyration’, a metric depicting the average distance of all atoms from a proteińs center of mass [151]. When *Reduced distance* value exceeds 1 (0.8 for min-max normalized values), it suggests that a residue is located further from the center than the average. This may influence its function and interaction within the protein‘s overarching structure [150].

*Shape and Surface* (n=3) gauges relationships between the physical form of an amino acid and its solvent exposure in folded proteins. Two measures describe the rate at which the accessible surface increases relative to the protein core distance [152]. Also included is the typical torsion angle a side chain adopts within folded proteins [153]. While high for small (glycine > alanine > proline) and polar amin residues (*e.g.*, serine), these measures are low for cysteine and methionine, reflecting their tight packing within globular proteins [152,153].

*Steric parameter* (n=6) describes measures regarding the steric complexity of an amino acid side chain, such as branching, symmetry, or side chain angles. Low for glycine and high for isoleucine, these factors can influence how a residue fits into protein structures and its interactions with adjacent residues [125].

#### 3.2.7 Structure-Activity

The ‘Structure-Activity’ category, the second smallest category with 31 scales in 6 subcategories, encapsulates the spectrum of structural dynamics from flexibility to stability. Flexibility, often associated with surface residues and hydrophilicity, is key for interactions at sites such as catalytic centers, binding domains, or antigenic regions [25,26]. Conversely, structural stability is fundamental to protein folding and is generally high in buried, hydrophobic residues.

*Flexibility* (n=11) refers to local residue movement within a protein (*e.g.*, positional changes or involvement in bending or twisting), comprising side chain and backbone flexibility [154]. Typically quantified by the B-factor in X-ray crystallography [155], it reflects structural flexibility in form of atomic fluctuation and thermal vibrations. While side chain flexibility does not imply backbone flexibility, both types can contribute to conformational changes, allowing flexibility in disordered and ordered regions [156]. Side chain flexibility enhances molecular interactions such as protein-ligand binding [157], whereas backbone flexibility is found in surface regions or epitopes [158]. Structure-breaking residues (proline, glycine), lysine, aspartic acid, and serine exhibit high flexibility [158–161]. *Flexibility (2 rigid neighbors)* (n=3) assesses the flexibility of a residue placed between two rigid neighboring residues, providing a context-specific view on structural flexibility. Under such conditions, lysine retains high flexibility, while the flexibility of aspartic acid, proline, serine moderately decreases, and that of glycine markedly drops [158,160].

*Stability* (n=7) denotes residueś contribution to enhancing protein stability. In contrast to *Flexibility* reflecting local movement, stability is a global property of entire proteins governed by intermolecular forces, such as hydrophobic interactions. Typically, buried residues increase stability by forming β-sheets, measured, for example, by the β-coil equilibrium [162], describing the preference of residues for extended over disordered coil structures. Prevalent residues include branched-chain (isoleucine, valine > leucine) and aromatic amino acids [62,162,163].

*Stability (helix-coil)* (n=4) quantifies the stability of residues using the helix-coil equilibrium. Higher scores indicate a preference for α-helices over disordered coil structures. Hence, residues common in α-helices, such as leucine and methionine, show higher *Stability (helix-coil)* scores. In contrast, residues more prevalent in β-sheets (*e.g.*, isoleucine and valine), tend to score lower in *Stability (helix-coil)* compared to *Stability* [162,164].

*Backbone-dynamics (-NH)* (n=2) reflects the mobility of a residués α-NH hydrogen atom within a polypeptide backbone, with higher values indicating backbone stability [165]. It is determined by α-NH chemical shifts using NMR, comparing hydrogen atom mobility in structured polypeptide to in random coil (highly flexible). High scores imply α-NH backbone stability, while low scores, typical in proline, indicate high mobility. Using an alternate method, spin-spin coupling, methionine and tryptophan score 0 [166].

*Backbone-dynamics (-CH)* (n=3) gauges the mobility of a residués α-CH hydrogen atom within a polypeptide, reflecting backbone stability [165]. These measures use α-CH chemical shifts to compare the hydrogen atom’s mobility between a structured backbone and a random coil. While tyrosine, asparagine, and lysine have high scores, low scores can be seen for valine, isoleucine, and especially glycine [166,167].

#### 3.2.8 Others

The ‘Others’ category, comprising 17 scales and 6 subcategories, contains scales which could not be reasonably assigned to other categories. It includes mutability and 5 groups of scales derived by principal component analysis (PCA). Essentially, PCA is a statistical technique that transforms a large set of correlated variables (*e.g.*, scales) into a smaller set of uncorrelated variables, called principal components (PC). These PCs encapsulate the maximum variance, thereby offering a succinct representation of the initial data and reducing the number of variables greatly. Moreover, it provides insight into the relative importance of these variables. This approach was employed by Sneath on 134 amino acid scales [168], yielding four principal components (PC1–PC4), which we utilized for our subcategory naming.

*Mutability* (n=3) refers to the likelihood of a residue to undergo mutations, calculated as the ratio of observed changes to its frequency of occurrence. This offers valuable insights into the evolutionary pressure exerted on specific residues within protein sequences [169].

*PC 1* (n=1), termed ‘aliphaticity’ and ‘typicalness’ by Sneath, represents mainly a residués aliphatic properties (*i.e.*, signifying the presence of linear, non-aromatic carbon chains) and its general prevalence in proteins [168].

*PC 2* (n=2), labeled ‘hydrogenation’ by Sneath, approximately corresponds to the inverse of the reactive group count in a residue, with lower values indicating higher reactivity. While proline and glycine (in Sneath’s PCA) have the highest values, acidic residues exhibit the lowest [168].

*PC 3* (n=2), called ‘aromaticity’ by Sneath, denotes the aromatic properties of residues, elevated not only in aromatic residues but also residues with cyclic side chain such as histidine [168].

*PC 4* (n=2), referred to as ‘hydroxythiolation’ and an ambiguous property by Sneath, potentially represent the ability of residues to form hydrogen bonds, prevalent in amino acids such as cysteine or serine [168].

*PC 5* (n=2) is another principal component vector derived by Wold et al [170], but the precise properties it encapsulates are not explicitly detailed.

### 3.2 Relations of scale subcategories

Understanding the relationships between scale subcategories is crucial for interpretation of machine learning results. We assessed the relations between scale subcategories using hierarchical clustering, correlation analysis, and PCA. To make subcategories comparable, they were represented by their average scales (see ‘2.3 Representation of scale subcategories by average scales’).

#### Hierarchical clustering of scale subcategories and amino acids

To explore the relations between all 67 subcategories, we hierarchically clustered their representative average scales based on their Euclidean distance using agglomerative clustering with complete distance [10,11] (**Fig. 13)**.

**Fig. 12.**
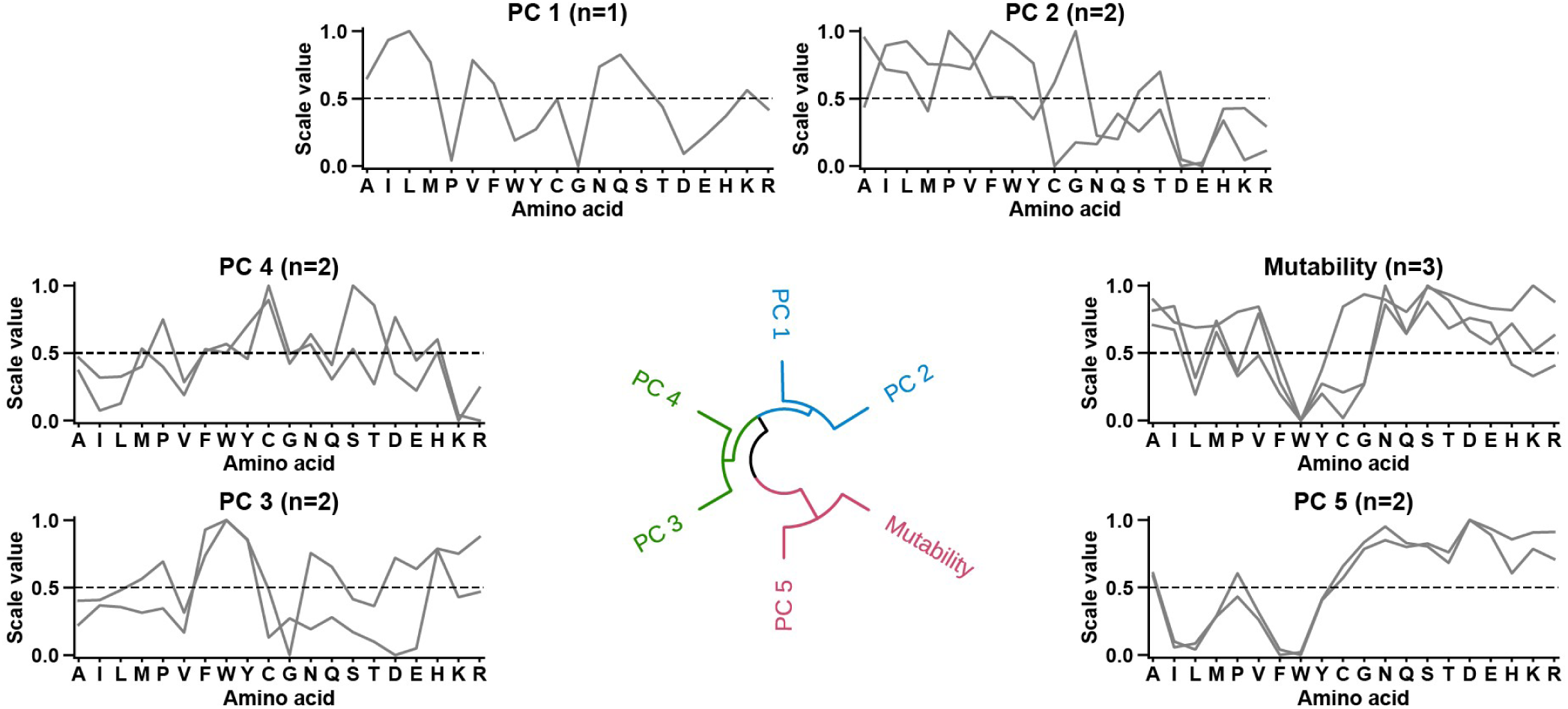
Others category. Circular dendrogram showing hierarchical clustering of all ‘Others’ subcategories. See **Fig. 5** for details.

**Fig. 13.**
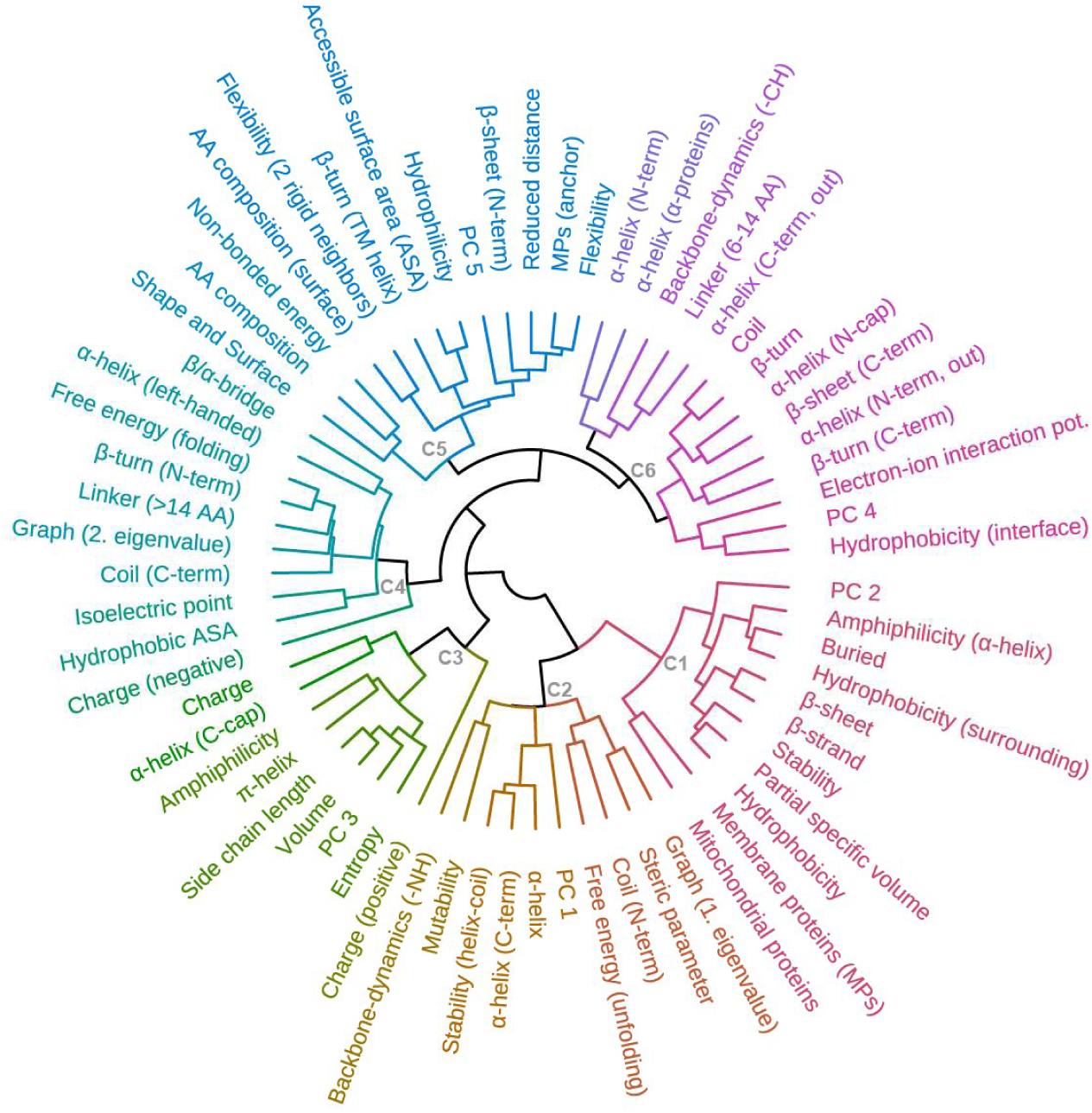
Hierarchical clustering of scale subcategories and amino acids. **a**, Circular dendrogram showing the following 6 clusters (C1–C6) of scale subcategories: C1 (*e.g.*, *Hydrophobicity* and *β-strand*), C2 (*e.g.*, *Steric parameter* and *α-helix*), C3 (*e.g.*, *Charge* and *α-helix (C-cap)*), C4 (*e.g.*, *Hydrophobic ASA* and *Linker (> 14 AA)*), C5 (*e.g.*, *Hydrophilicity* and *Flexibility*), and C6 (*e.g.*, *Coil* and *β-turn*).

The 6 following clusters (C1–C6) were found: C1 reveals that hydrophobicity correlates with stability, frequently seen in buried residues, and driving the formation β-strands and β-sheets; C2 links α-helix propensities to conformational stability (*Free energy (unfolding)*) and steric complexity; C3 highlights the correlation between entropy, volume, side chain length, and charge, all characteristic in C-terminal residues of α-helices; C4 relates hydrophobic ASA, long linkers, and conformational instability (*Free energy (folding)*)—pronounced for proline; C5 demonstrates hydrophilicity relating to ASA, flexibility, and membrane anchor or surface residues; and C6 underlines a close conformational link between propensities to form coils, β-turns, and medium-sized linkers. Here, the connection between *Coil* and α-*helix (N-cap)* underscores that the α-helical N-terminus is less stable than the C-terminus [171], while the association of *β-turn* with *β-sheet (C-term)* illustrates the process of anti-parallel β-sheet formation, characterized by β-strands that end at the C-terminus and sharp backbone reversal via a β-turn [108,172].

In essence, C1–C2 represent core-forming properties, especially α-helix and β-sheet propensities. C3–C4 pertain to conformational transitions and molecular interactions, while C5 highlights surface-related properties. Finally, C6 underscores the contribution of coils and β-turns in terminating structured conformations.

#### Correlation analysis of subcategories

Scale-based machine learning can identify features not anticipated or seemingly inconsistent with our biological understanding. To enhance their interpretation, we investigated correlations and anti-correlations between average scales of subcategories (**Fig. 14**).

**Fig. 14.**
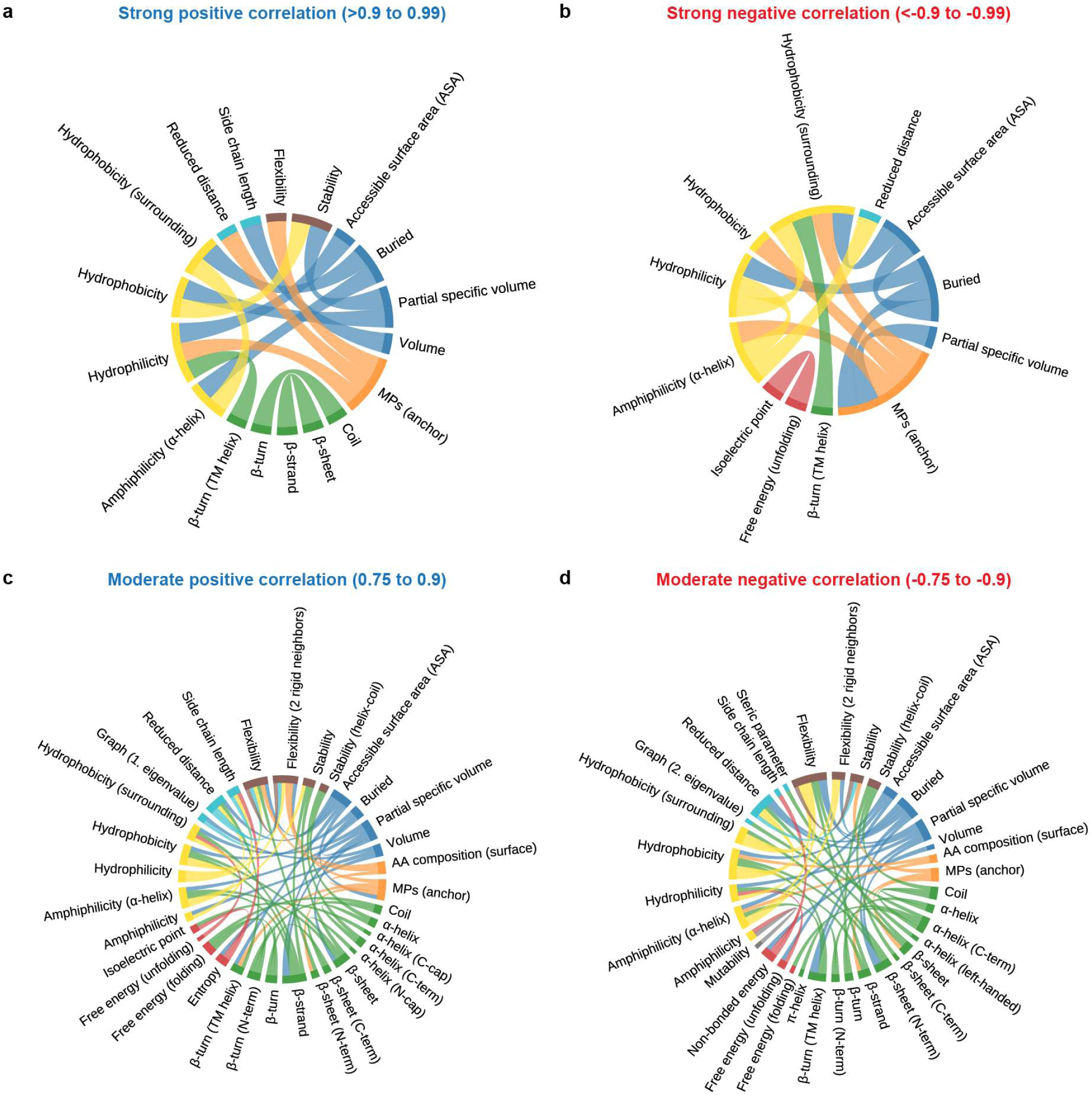
Correlations between average scales across scale categories. Chord diagrams showing the correlation and anti-correlation of subcategories filtered for ranges of Pearson correlation. Connections are formed between subcategories when their correlation lies within the indicated correlation range. Colors indicate the category to which the subcategories are assigned: ASA/Volume (blue), Composition (orange), Conformation (green), Energy (red), Polarity (yellow), Others (gray), Shape (light blue), and Structure-Activity (brown). Subcategories with fewer than 3 scales, such as those related to charge, were omitted for clarity. Self-referencing correlation of 1 or - 1 were disregarded.

Among strongly positively correlated subcategories (Pearson’s *r*>0.9 to 0.99, **Fig. 14a**), three groups underline the connection between polarity and protein folding: (a) *Hydrophobicity*, *Partial specific volume*, and *Stability*, (b) *Hydrophilicity, Accessible surface area (ASA)*, and *MPs (anchor)*, which is additionally highly correlated with *Flexibility* and *Reduced distance*; and (c) *Amphiphilicity (α-helix)*, *Buried*, and *Hydrophobicity (surrounding)*. Other notable highly correlated pairs include *Volume*/*Side chain length*, *AA composition*/*MPs (single-spanning)*, and the conformation pairs *Coil*/*β-turn* and *β-sheet*/*β-strand*.

Strong negative correlations (Pearson’s *r*<-0.9 to -0.99, **Fig. 14b**) shed new light on three polarity groups: While *Hydrophobicity* anti-correlates with *MPs (anchor)*, *Hydrophilicity* anti-correlates with *Amphiphilicity (α-helix)*, *Hydrophobicity (surrounding)*, and *Buried*. In addition, the *Accessible surface area (ASA)*/*Buried* anti-correlation highlights the clear distinction between surface and buried residues. The anti-correlation of *Isoelectric point* and *Free energy (unfolding)* (conformational stability) shows that basic residues can potentially decrease stability due to the disruptive potential of their positive charge when buried, albeit context-dependent. Notably, *MPs (anchor)* inversely correlates with five other subcategories including *Buried* and *Amphiphilicity (α helix)*, underlaying its unique residue signature, dominated by proline, aspartic acid, and lysine.

Moderate positive correlations (Pearson’s *r*=0.75 to 0.9, **Fig. 14c**) reveal a complex picture, especially for three major conformations: (a) *Coil* correlates with *Free energy (folding)* (conformational instability) and various β-turn subcategories; (b) *α-helix* correlates with *α-helix (C-term)* and *Stability (helix-coil)*, while *α-helix (C-cap)* correlates with *Isoelectric point* (basicity); (c) Extended conformations (*β-sheet*, *β-strand*) correlate with *Hydrophobicity*, *Stability* and *Partial specific volume*. Interestingly, structure termination (*β-sheet (C-term)*, *α-helix (N-cap*) relates to *β-turn*, which highlights their structural overlapping. The correlation of *Entropy* with *Volume* and *Side chain length* indicates that larger residues contribute to structural variety (α-helix or β-sheet), unlike the helix-breaking proline or the helix-forming alanine. *Flexibility* shows broad correlations, including *Hydrophilicity* and *Accessible surface area (ASA)*.

We also identified diverse moderate negative correlations (Pearson’s *r*=-0.75 to -0.9, **Fig. 14d**). As expected, *Coil* anti-correlates with *α-helix*, as do extended conformations with *Hydrophilicity* and *Flexibility*. Notably, *Amphiphilicity* is negatively correlated with *Free energy unfolding* and *Mutability*, suggesting that amphiphilic residues can be less stable (*e.g.*, arginine) and less mutation-prone (*e.g.*, tryptophan). *Non-bonded energy* inversely correlates with *Volume* and *Side chain length*, showing that small residues have reduced van der Waals forces and electrostatic interactions. See **Supplementary Table 4** for detailed results of our correlation analysis.

#### Principal component analysis of scale subcategories

PCA provides another way to visualize the relationships between scale subcategories. To obtain a two-dimensional ‘snapshot’ of these relationships, we divided the scale subcategories into four roughly equally sized sets from different categories, plotting the first two principal components for each (**Fig. 15a**).

**Fig. 15.**
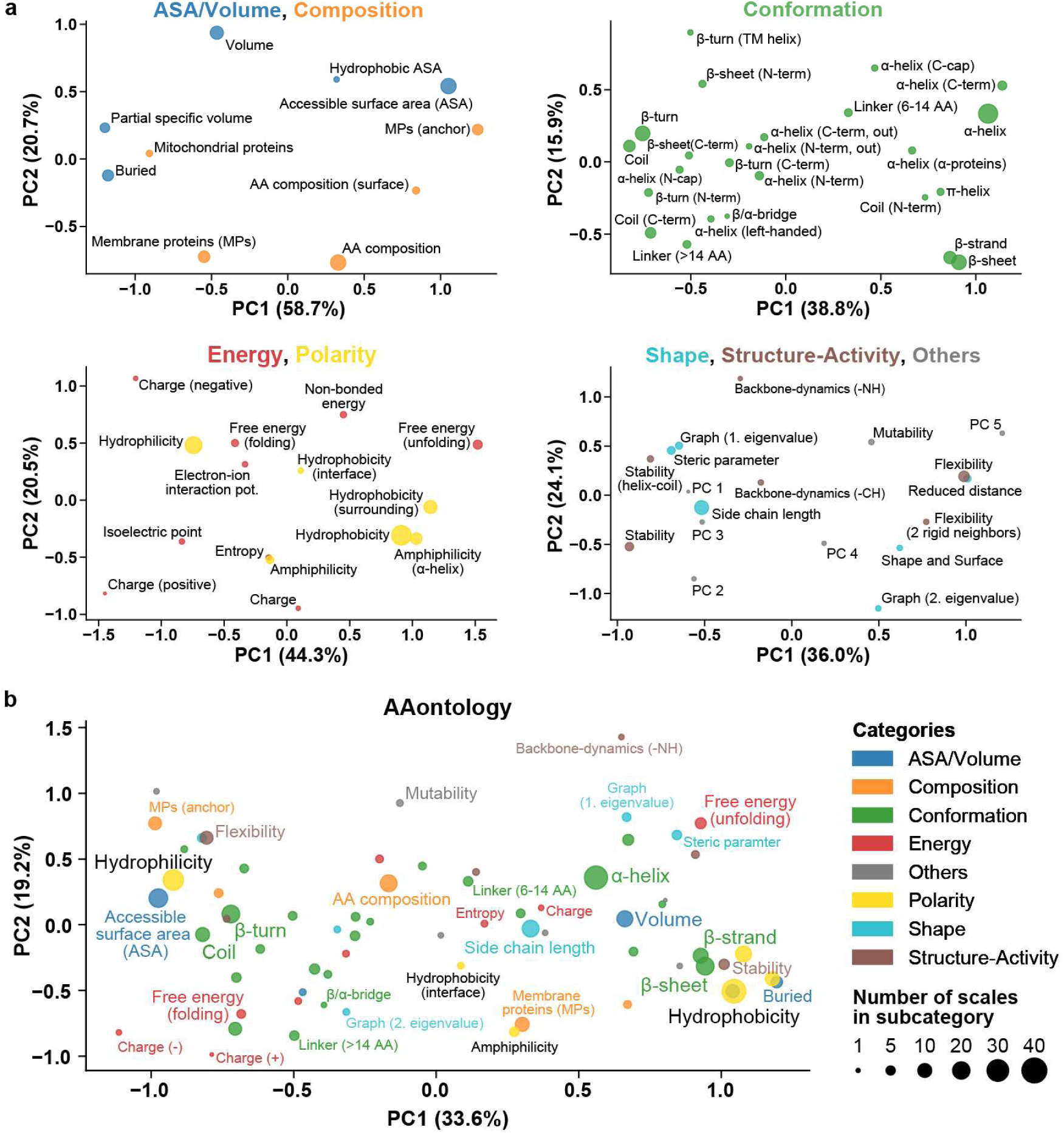
Relations between subcategories. Principal component (PC) analysis of average scales representing scale subcategories. **a**, Four separate PCA for subcategories of following sets of categories: ASA/Volume and Composition; Conformation; Energy and Polarity; Shape, Structure-Activity, and Others. **b**, PCA for all subcategories contained in AAontology with important subcategories being annotated. Each points represents a subcategory, color-coded by category; and their size is indicating the number of scales that are assigned to the subcategory.

Our first PCA involved ‘ASA/Volume’ and ‘Composition’ subcategories. The first dimension, PC1 (56.2% variance explained), reveals a spectrum between buried and solvent accessible surface, relating *Buried* with *Mitochondrial Proteins*, and *Accessible surface area (ASA)* with *MPs (anchor)*. The second dimension, PC2 (23.3%), differentiates *Volume* from *Membrane proteins (MP)* and *AA composition*, implying that larger residues occur less frequently, especially in membrane proteins.

‘Conformation’ subcategories were analyzed together due to their large number. The first PC (38.8%) distinguishes unstructured (*Coil*, *Linker (> 14 AA)*) and *β-turn* from extended (*β-strand*, *β-sheet*) and α-helical conformations, which are both separated along the second PC (15.9%). Interestingly, subcategories representing termination of β-sheets or α-helices are closely related to *Coil* and *β-turn*.

We grouped ‘Energy’ and ‘Polarity’ subcategories since hydrophobicity is often measured as transfer free energy from a non-polar solvent to water. Polarity (hydrophilicity vs. hydrophobicity) is reflected by the first PC (43.6%), relating *Free energy (folding)* to *Hydrophilicity* and *Free energy (unfolding)* to *Hydrophobicity*, reinforcing the polarity-conformational stability relationship. Interestingly, *Entropy* closely aligns with *Amphiphilicity*. The second PC (21.4%) differentiates positive from negative charge. We also analyzed ‘Shape’, ‘Structure-Activity’, and ‘Others’ categories. The first PC (36.0%) underscores the *Stability*-*Flexibility* dichotomy, relating *Stability* to steric complexity (*e.g.*, *Steric parameter* and *Side chain length*), and *Flexibility* to *Reduced distance*. The second PC (23.1%) separates orthogonal aspects of a residue’s structural graph representation. embodied by *Graph (1. eigenvalue)* and *Graph (2. eigenvalue)*.

In a comprehensive PCA of all subcategories (**Fig. 15b**), the polarity spectrum is represented in the PC1 (33.4%). *Hydrophilicity* relates to *Accessible surface area (ASA)*, *Flexibility*, *Coil*, and *β-turn*, while *Hydrophobicity* links with *Buried*, *Stability*, and structured conformations—*β-strand* and *β-sheet* more than *α-helix*. These ordered conformations are separated along the PC2 (19.1%), which also distinguishes the two complementary graph-theoretic measures of *Graph (1. eigenvalue)* and *Graph (2. eigenvalue)*. Both PCs together discriminate *Free energy (unfolding)* from charge (negative and positive) and *Free energy (folding*).

Overall, this analysis elucidates the interplay between higher-order physicochemical properties in protein folding, primarily driven by the hydrophilicity-hydrophobicity dichotomy. Protein folding is a two-phase process orchestrated by fundamental intermolecular forces (highlighted in *italic lowercase*). The first phase comprises the formation of secondary structures through a balance of attractive (mainly backbone *hydrogen bonding* and also *hydrophobic interactions*) and repulsive forces (*electrostatic repulsion* and *steric restrictions*) [173], reflected in the ‘ASA/Volume’, ‘Energy’, ‘Polarity’, and ‘Structure-Activity’ categories. The subsequent phase involves inter-amino acid forces, captured by the ‘Polarity’ and ‘Energy’ categories. Polar residues participate in diverse interactions (*dipole-dipole* or *ion-dipole interactions*, *hydrogen bonds*, or *ionic interactions*), while non-polar residues engage in weaker *van der Waals* and *hydrophobic interactions*. The latter drives hydrophobic clustering to minimize water exposure, known as the hydrophobic effect, which is crucial for protein stability [174,175]. Protein structures emerge from this interplay of intermolecular forces, influenced further by the molecular and cellular context.

## 4. Discussion

### Advantages and challenges for AAontology

AAontology, with its two-level classification and in-depth subcategory analysis, enables a broader understanding of amino acid properties and provides a rigorous interpretation for scale-based machine learning. AAontology derives its strength from consensus-driven, robustly explained subcategories. Furthermore, it is a transparent, modular, and consistent framework to incorporate new amino acids properties. Its versatile nature permits applications across a wide range of computational biology problems, such as mutation analysis or protein design.

The abundance of property scales collated over six decades holds potential pitfalls. Specifically, issues may arise due to incomplete knowledge and the quality of the scales included. While AAontology helps to structure existing knowledge about amino acid properties, it is inherently limited by our current understanding of these characteristics. Any error or bias may inadvertently be carried over into computational approaches based on AAontology.

To tackle these pitfalls, scales can be revised or updated, as was done for CHAM830107 from the *Charge (negative)* subcategory and polarity scales, respectively. Furthermore, new subcategories could be defined, reflecting, for example, different types of β-strands [107] or participation in certain catalytic processes such as nucleophilic attacks (serine, cysteine, and histidine) or transition state stabilization by aromatic rings (*e.g.*, tyrosine) or charge (*e.g.*, aspartic acid or lysine).

A broader concern arises considering the general limitations of scale-based machine learning models. Their performance varies considerably depending on the prediction task, chosen model, and scale dataset used [48]. For instance, simple randomly created scales or one-hot encodings of residues have outperformed biologically meaningful property scales, however, only for a specific redundancy-reduced scale set [176]. Additionally, the array of scales and subcategories in AAontology could add complexity, challenging novice researchers.

Overcoming these limitations could involve knowledge-based pre-selection of categories from AAontology, coupled with the selection of redundancy-reduced scale sets via our AAclust framework. Another solution is offered by large protein language models, overpassing simple scale-based approaches [177–179], and thereby propelling the field of protein prediction [180].

### Do protein embeddings abolish scale-based machine learning?

Scale-based approaches (since the 1970s [80,112]) and Multiple Sequence Alignment (MSA)-based models (since the late 1980s [181–183]), leveraging evolutionary sequence information, have long been instrumental in protein structure and function prediction. However, the early 2010s saw the advent of deep learning models including Convolutional Neural Networks (CNNs) [184,185] and Long Short-Term Memory (LSTMs) [186]. In the late 2010s, transformer models [187] emerged, combining CNN local structure capture and LSTM long-range dependency handling, which lead to breakthroughs such as AlphaFold [188] in protein structure prediction.

Recent state-of-the-art protein language models, such as ProtT5 [189,190], utilize transformer architectures to improve protein prediction tasks. These models, trained on billions of sequences, aim to decipher the ‘language of life’ [189,191] encoded by proteins. To this end, they produce a self-learned amino acid representation, called protein embeddings, that can be viewed as property scales enriched by both nearby and distant sequence context. Beside their superior prediction performance, these protein embeddings are advantageous in their simple application and unbiased nature because they are alignment-free [190] and do not rely on hand-crafted features [179]. They can also serve as input to other machine learning models via transfer learning [192], enabling their application to moderately-sized datasets.

A significant drawback is, however, the lack of interpretability of protein embeddings [8,9], as they are an opaque numerical representation of proteins. This challenge could potentially be addressed by AAontology. Mapping protein embeddings onto scale subcategories could merge the predictive power of embeddings with the interpretability provided by AAontology. Improved interpretability would enhance the biological insight derived from protein embeddings. The process of mapping might even expose yet unknown properties [193], that one could call the ‘dark matter’ of protein biology, in an analogy previously used in this field [194,195]. Consequently, we foresee a synergistic relationship between AAontology and protein embeddings, fostering a novel understanding of proteins.

### AAontology as the fundament to understand protein biology

AAontology offers a framework to study how amino acids determine protein structure and function. While AlphaFold can produce accurate snapshots of protein structures, simulating complex and/or slow molecular processes remains still elusive [196]. AAontology can fill this gap by providing a deep understanding of amino acid properties to elucidate molecular mechanisms such as substrate or epitope recognition.

Additionally, AAontology facilitates a multi-dimensional understanding of amino acid changes, especially pertaining to disease-related mutations [35–40]. For instance, an alanine-to-lysine substitution not only introduces a positive charge and increases volume and hydrophilicity, but also increases the hydrophobic ASA, entropy, and flexibility within two rigid neighbors, while decreasing buriability, certain shape and surface properties and the propensity to form amphiphilic α-helices.

Moreover, we envision that AAontology serves as a cornerstone in linking physicochemical properties to protein functions and dysfunctions—as for example done for post-translational modifications [197]. Notable examples of connections to protein functions include (a) substrate cleavage facilitated by the formation of extended β-strands [198–201]; (b) epitope recognition influenced by hydrophobicity and hydrogen bonds [202,203]; and (c) protein-protein interactions [204,205] (*e.g.*, chaperone-client interactions [206,207]), protein-peptide interactions (involving peptides with extended structures, but also α-helix or β-turn conformation [105]), and protein interactions with other biomolecules such as DNA [208], RNA [208], lipids [208], or small molecules [209]. Physiochemical properties are further important for subcellular localization [21], protein stability [210], or protein trafficking [211], next to general cellular functions such as signaling [25,26] or cell division [212].

Protein aggregation is a crucial example for protein dysfunction. Triggered by increased hydrophobicity and β-sheet tendency, protein aggregation is associated with over 40 human diseases including various neurodegenerative disease (*e.g.*, Alzheimer’s, Parkinson, Huntington, and prion disease) and type II diabetes [32–34]. Numerous efforts have been made to predict aggregation-prone sequence regions [24,213,214], yet they still need to be refined. Oncogenicity, another key example, is associated with altered protein folding due to changes in hydrophobicity, charge, or conformational characteristics [29–31]. Other examples encompass alterations in hydrophobicity leading to cystic fibrosis [215], loss of charge in voltage-gated sodium channel associated with epilepsy [216], or decreases α-helicity in myosin causing cardiomyopathy [217]. A systematic overview of the relationships between physicochemical properties and protein functions and dysfunction could lay the groundwork for a deeper understanding of protein biology. This would further foster functional annotations [218–220] of novel or so far poorly characterized proteins.

Overall, AAontology, in conjunction with tools like AlphaFold or ProtT5, is expected to unravel molecular mechanisms and unfold a systematic understanding of protein functions. This will augment numerous applications that rely on knowledge-based decisions. These include the development of biocompatible materials (*e.g.*, amino acid-based surfactants (*i.e.*, surface active agents) [221,222]), rational protein engineering [193] (*e.g.*, enzyme engineering [223]), and drug design of peptides [224,225] or proteins [210,226], such as vaccines [227,228] or, especially, antibodies [229–232].

## 5. Conclusion

Through a semi-automatic process involving clustering and manual refinement, we classified 586 amino acid scales into 8 categories and 67 subcategories, resulting in AAontology. This two-level classification also provides a detailed subcategory description and an extensive analysis of their relationships, bolstering interpretable machine learning models for protein bioinformatics. Beyond hierarchical clustering, PCA allowed to map these subcategories onto the major physicochemical spectra of polarity (hydrophobicity vs hydrophilicity) and conformational stability, thereby painting a holistic picture of the properties governing life.

AAontology, when combined with our AAclust framework, could overcome limitations of scale-based machine learning in protein prediction. Our plans include integrating AAontology with protein embeddings, such as ProtT5, enriching high predictive power with interpretability and possibly unveiling hitherto unknown amino acid properties. Complementing tools such as AlphaFold, AAontology could foster the deciphering of molecular mechanisms encapsulated in protein structures. Linking physicochemical properties and protein function/dysfunction, AAontology will be a useful decision support tool for mutation analysis and drug design.

## Author Contributions

S.B. conceived the study, performed all computational analysis, conducted the manual refinement, and wrote the manuscript. F.K., H.S., and D.F. provided substantial support through critical discussions and participated in manuscript editing.

## Acknowledgements

We thank Dieter Langosch for engaging in critical discussions regarding physicochemical properties, particularly on aspects related to polarity and flexibility.

## Code availability

AAontology is a foundational component of AAanalysis, a Python-based framework for interpretable protein prediction, which will be made available in a forthcoming publication.

## Supplementary material

Supplementary Table 1–4, including all 586 amino acid scales and the AAontology scale classification, can be found online with this manuscript.

